# Cortical and Hippocampal Pathways to Mental Continuity

**DOI:** 10.64898/2026.09.04.749493

**Authors:** Xian Li, Hongmi Lee, Christopher Honey

## Abstract

How do people preserve mental continuity when tasks are interrupted? One established pathway runs through the hippocampus, which stores snapshots of our mental context for later reinstatement. We propose a second pathway: the brain’s default network sustains the interrupted situation as a background process while other processing continues. In 73 adults listening to interrupted stories during functional brain imaging, the posterior medial cortex sustained a reliable and selective representation of the stages of the interrupted narrative. This background representation changed gradually over narrative time and persisted in a transformed format through silence and through a demanding secondary task. Its persistence predicted how reliably the default network resumed the story and how well listeners later recalled it, revealing complementary hippocampal and cortical pathways to mental continuity.

**One-Sentence Summary:** Posterior medial cortex carries narrative context through interruption in transformed form, predicting resumption and memory

---

While reading a book we are interrupted by a phone message. It takes thirty seconds to answer the message, then resume where we left off. Interruptions of this kind are commonplace: conversations are diverted, plans are set aside, and books are put down. However, for human thought and behavior to extend over minutes, we must sustain mental context beyond such interruptions (*1*). What is happening in the brain during those thirty seconds, and why is our mental context better preserved on some occasions than others?

One pathway to mental continuity relies on discrete storage and retrieval. We can store a trace of high-level mental context at the start of an interruption, via the hippocampus, and then retrieve that mental context when the interruption ends. However, there is reason to believe in a second pathway for maintaining high level mental context. First, even if long-term retrieval is employed, it requires a cue or index of the past context, and this would need to survive during interruption.

Second, people commonly experience that recent high-level context persists in the back of their minds, without volition, even when they switch to a new task (*2, 3*). Third, the speed and automaticity of recovering from reading interruptions is difficult to reconcile with an un-cued search and retrieval from long term memory (*4*). For these reasons, we hypothesized that, in addition to the hippocampal pathway, there is a neocortical pathway for situational context to persist in the background of interrupting tasks.

The human default mode network (DMN) is a candidate substrate for maintaining situational context: it is implicated in narrative comprehension and immersion, event structure, and internally generated representations, and its activity tracks information over long timescales during naturalistic cognition (*5–10*). Indeed, DMN regions can integrate narrative information over tens of seconds even when the hippocampus is damaged (*11*). We therefore hypothesized that the “background” representation is maintained within the DMN.

If DMN circuits maintain a background representation of mental context, what format should it take? Early neuroscience studies of working memory identified “delay-period activity” in lateral prefrontal neurons which sustained the representation of a stimulus or task (*12*), while more recent studies have argued that information can be maintained in an orthogonal (*13*) or even inverted format (*14, 15*) in frontoparietal or sensory circuits. If background context is automatically maintained in DMN circuits similarly to these working memory traces, then DMN traces may also adopt a “sustained” delay period signature or a transformed format.

Using functional magnetic resonance imaging (fMRI) during extended narratives with controlled interruptions, we sought to determine which brain systems maintain an interruption-resistant background representation of narrative context. We first measured whether interruption patterns were reliable across participants and selective to the narrative context, then whether they changed gradually over narrative time and persisted when interruption epochs were filled with a theory-of-mind task. We found many of these signatures within DMN regions of the human brain, but only the posterior medial cortex (PMC) expressed all of them. Examining the representations in PMC, we found that they did not simply sustain the story-phase pattern: PMC carried the interrupted narrative in a rotated format. Moreover, the persistence of this transformed representation predicted the resumption of the narrative neural state (when the interruption ended) as well as participant’s recall accuracy for the narrative. These findings suggest a neural mechanism for mental continuity: contextual representations in PMC can be transformed into a protected background pattern.

## Narrative processing with and without interruptions

Seventy-three participants listened to a 14-min story during fMRI in one of four conditions (materials and methods): the intact story continuously presented (Continuous, CT; n = 16), the intact story interrupted by silent pauses (Intact-Pause, IP; n = 19), the intact story interrupted by a story-irrelevant theory-of-mind task (Intact-ToM, IT; n = 19), or a scrambled version of the story interrupted by silent pauses (Scram-Pause, SP; n = 19). The story was divided into 18 segments of 30-60 s, with 17 intervening interruption epochs of 20-30 s in the interrupted conditions. Participants verbally recalled the story afterwards. A subset of participants additionally completed behavioral tests and surveys on story comprehension, memory, and thoughts outside the scanner.

Participants processed the intact narratives successfully despite interruption. Recall and comprehension of the story were preserved in both interrupted intact-story conditions relative to the continuous baseline (*16*), while scrambling impaired performance. Recall differed across conditions [one-way ANOVA, *F*(3, 67) = 5.95, *P* = 0.001], with lower recall in SP than CT (*P* = 0.015) and IP (P < 0.001; Bonferroni-corrected Welch t tests here and below). Comprehension also differed across conditions [*F*(3, 55) = 8.01, *P* < 0.001], with lower comprehension in SP than CT (P < 0.001), IP (*P* = 0.002), and IT (*P* = 0.013).

We next examined the story-driven neural responses. Whole-brain inter-subject correlation (ISC) maps during story listening showed robust cross-participant synchrony spanning early auditory cortex and high-level cortical regions associated with narrative comprehension (*17*), including canonical default-mode regions under the IP condition (Fig. 1G) and other conditions (supplementary text S1, fig. S3). In addition, hippocampal activity increased at interruption onset [mean post-minus-pre difference = 0.049; t(56) = 3.20, P = 0.002; Fig. 1H], consistent with prior evidence that hippocampal responses at post-event periods and event boundaries support encoding of naturalistic episodes (*18, 19*). Whole brain boundary responses were consistent with prior literature (*20*; supplementary text S2).

**Fig 1.**
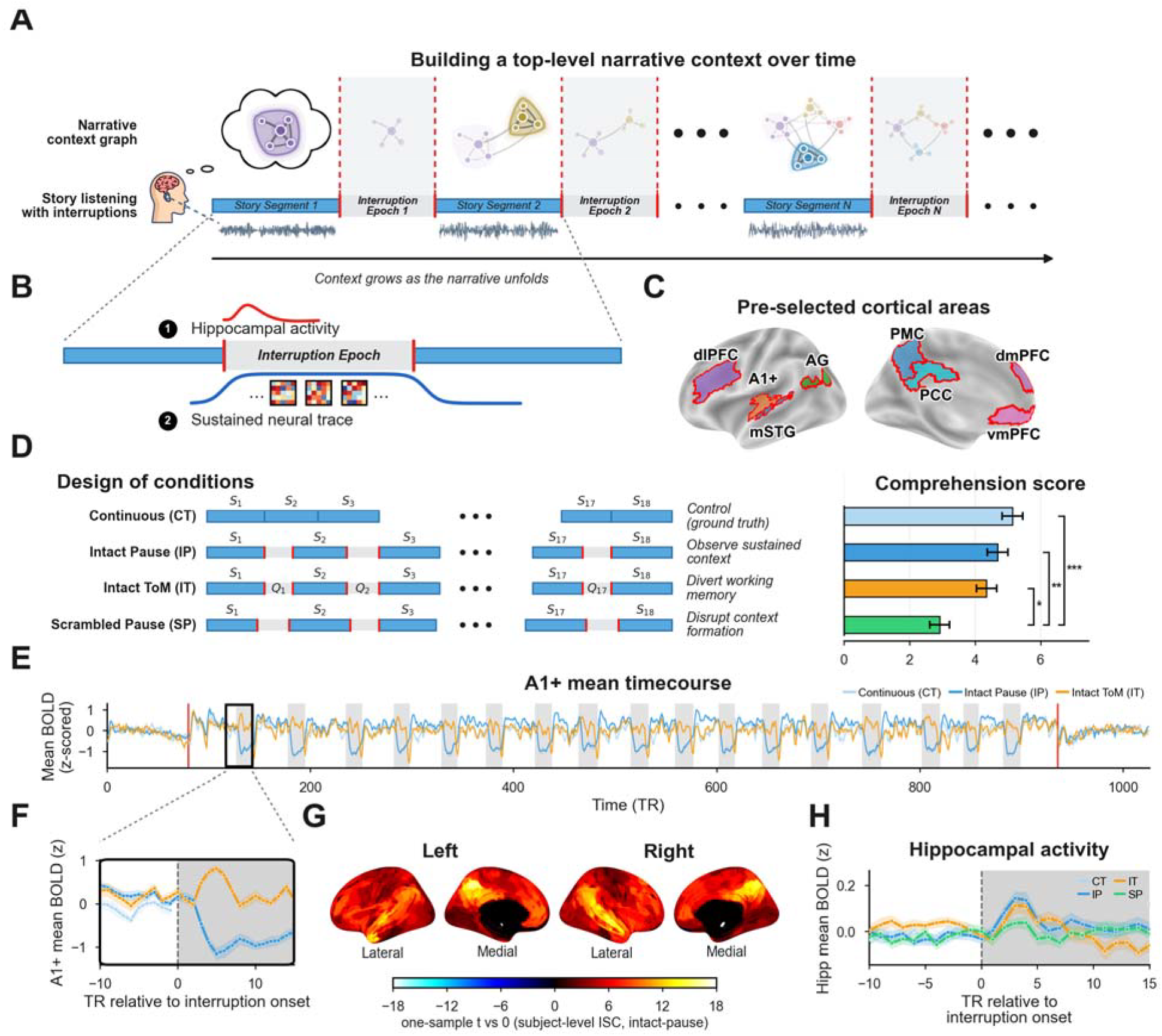
Study design, behavior, and story-driven neural responses. (A) Schematic of narrative context built over the story. As the story unfolds, each story segment’s event network (saturated colors) is integrated with accumulated past context (pale colors), progressively constructing the narrative context. Interruption epochs may sustain some format of this accumulated context. (B) Two candidate mechanisms for carrying context across an interruption: a hippocampal boundary response (1) and a sustained high-level neural trace (2). (C) Pre-selected cortical regions of interest. (D) The four experimental conditions and their associated story comprehension scores (mean ± SEM; Bonferroni-corrected Welch t tests: *P < 0.05, **P < 0.01, ***P < 0.001). (E) Mean z-scored A1+ signal across the full run for the continuous (CT), intact-pause (IP) and intact-ToM (IT) conditions; gray shaded bands mark interruption epochs. (F) Zoomed-in A1+ signal around the first interruption onset. (G) Whole-brain inter-subject correlation (ISC) during story listening in the IP group; the map shows the one-sample t statistic against zero across participant-level ISC maps (n = 19). (H) Hippocampal activity aligned to interruption onset (mean ± SEM per condition).

### Reliable, epoch-specific PMC interruption patterns evolved with the coherent narrative

We next tested whether interruption periods contained information about the interrupted narrative context. We analyzed multivoxel activity patterns during each interruption epoch across pre-selected default-mode, memory/control, auditory, and language regions (materials and methods, Regions of interest). For each region, we computed inter-subject pattern correlation (ISPC) against the group-mean pattern of the remaining participants and examined whether their interruption patterns were reliable across participants (supplementary text S3). We then asked whether the regions with reliable patterns were also selective to the preceding narrative epoch and organized across the unfolding story.

#### PMC showed reliable, epoch-specific interruption patterns

The PMC patterns were reliable across participants and more similar across participants for matching than mismatching epochs for both the IP [Fig. 2, A to C; selectivity = 0.034, P = 2 × 10^−4^, n = 19; permutation tests here and below unless named otherwise] and the SP group [selectivity = 0.015, P = 0.027], though SP was weaker than IP [Welch two-sided t(34.4) = 2.36, P = 0.024, d = 0.76]. Further, the epoch-selective interruption pattern in the IP group did not match that seen for the same epoch in the SP group (supplementary text S4). Thus, when the narrative scaffold was disrupted, a different and smaller amount of story-locked information persisted into the interruption epoch. Critically, when participants performed a story-irrelevant theory-of-mind task during interruption, their PMC patterns still matched the IP group in an epoch-specific manner [IT-IP: selectivity = 0.021, P = 0.003]. Thus, PMC interruption patterns retained information about the interrupted story segment even when working memory was occupied by a separate task.

**Fig 2.**
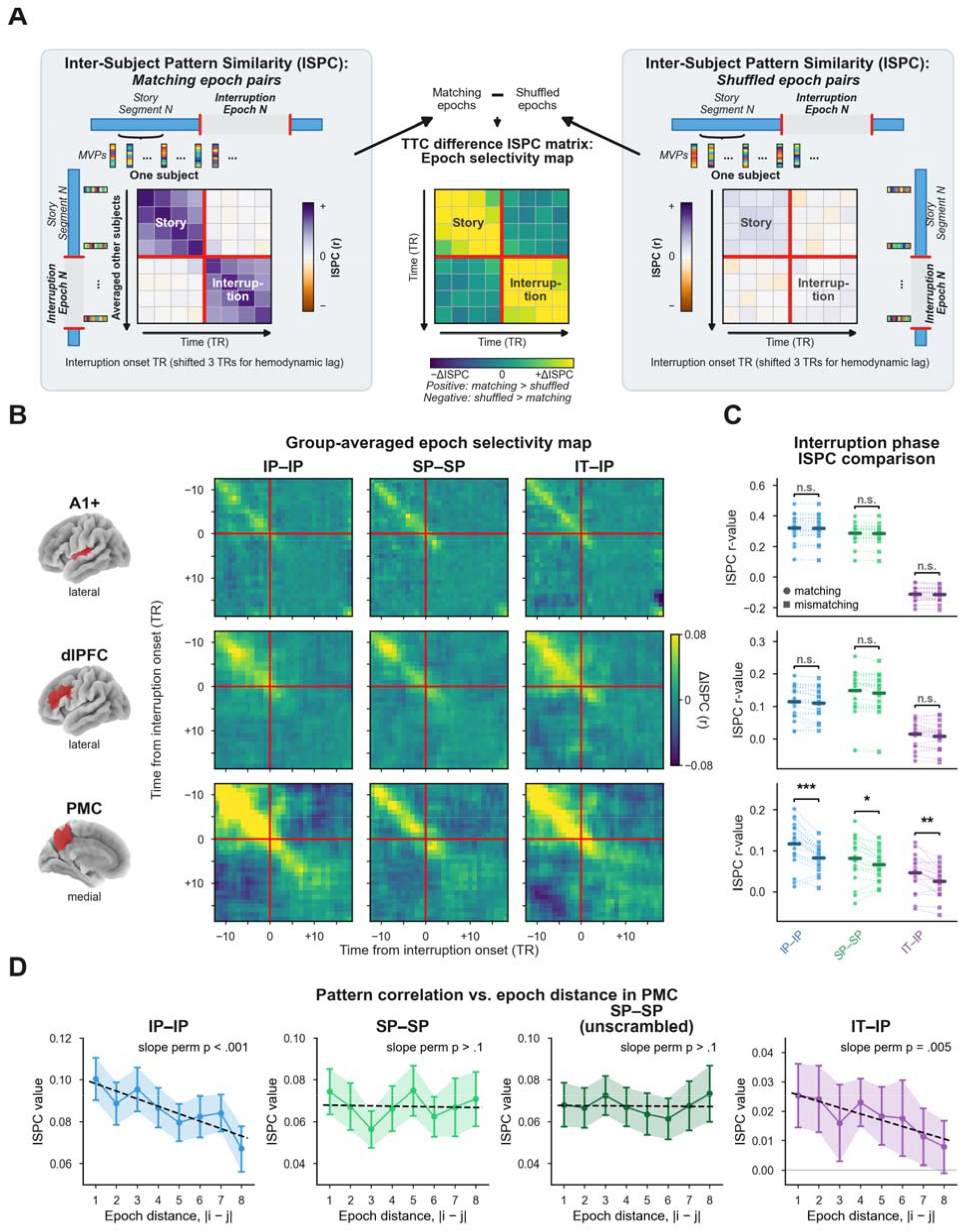
Reliable, epoch-specific interruption patterns that evolve with the narrative. (A) Time-by-time inter-subject pattern correlation (ISPC) analysis. Each participant’s multivoxel patterns around each interruption were correlated with group-mean patterns from matching versus mismatching narrative epochs. Epoch selectivity at each time-time pair is the difference (matching correlation − mismatching correlation); positive values indicate greater similarity for matching epochs. The labels IP-IP, SP-SP, and IT-IP denote the participant group and the comparison-template group, respectively. For the unscrambled SP-SP comparison, the pairwise distances of SP interruption epochs are defined according to distance in the intact narrative sequence. (B) Groupaveraged epoch-selectivity maps for A1+, dlPFC, and PMC under the IP-IP, SP-SP, and IT-IP comparisons (n = 19 each). (C) Interruption-phase ISPC for matching versus mismatching epochs. Horizontal bars indicate group means; dots, single participants. Epoch selectivity was assessed via within-participant label-shuffle permutation tests (10,000 iterations, one-sided); ***P < 0.001, **P < 0.01, *P < 0.05; n.s., not significant. (D) PMC interruption-phase ISPC as a function of narrative epoch distance (mean ± SEM) for IP-IP, SP-SP, unscrambled SP-SP, and IT-IP (n = 19 per scheme); group-mean slopes and their two-sided permutation P values are shown.

#### PMC interruption patterns evolved across the intact but not the scrambled narrative

The interruption patterns in the PMC changed gradually across the epochs of the intact narrative. The similarity between PMC patterns shared across the IP group decreased with epoch distance [Fig. 2D; IP-IP: distance slope b = −3.6 × 10^−3^, P = 9 × 10^−4^]. In contrast, the SP group showed no distance-dependent organization in the presented scrambled sequence [SP-SP: b = 1.3 × 10^−5^, P = 0.989] or when analyzed as an unscrambled sequence [SP-SP-unscrambled: b = 6.6 × 10^−5^, P = 0.943]. The IT-IP slope was also negative [IT-IP: b = −2.2 × 10^−3^, P = 0.005], so the gradual change persisted when the interruption was filled by the theory-of-mind task.

Across the full ROI set, several regions, including the angular gyrus, exhibited subsets of these story-locked signatures, but PMC was the only region showing the full pattern: reliable and epoch-specific interruption pattern, persistence under a competing ToM task, and gradual evolution across the intact but not scrambled narrative (supplementary text S5). The PMC signatures generalized to a second narrative (supplementary text S7), which identified PMC as the primary candidate substrate for an interruption-resistant background representation.

### Narrative context was sustained in a transformed format during interruption

We next investigated the format in which narrative information persisted. If the prior context were maintained as a conventional active representation, interruption-period patterns should resemble the preceding story-period patterns. Alternatively, PMC could preserve the interrupted context in a transformed format that could reduce interference with the interruption phase processing. We tested these alternatives by comparing multivoxel patterns from each story segment with patterns from the following interruption epoch.

#### PMC interruption patterns were transformed from their preceding story patterns

For each participant and epoch, we computed the similarity between that participant’s story-phase pattern and the average interruption-phase pattern measured across all other participants. Using ISPC between story- and interruption-window template patterns, each averaged over 10 TRs (repetition time, TR = 1.5 s), PMC showed reliable negative correlations from story to interruption phase in each condition [IP-IP: Fisher-z = −0.245, P < 1 × 10^−4^; SP-SP: −0.209, sign-flip P < 1 × 10^−4^; IT-IT: −0.171, P = 2 × 10^−4^; all n = 19; materials and methods, Story-to-interruption transformation analysis]; this negative correlation remained stable throughout the interruption window (Fig. 3A). The transformation was epoch-specific: matching story-interruption epochs were more negatively correlated than mismatched in all three interrupted conditions [Δ = matching − mismatching; IP-IP: Δ = −0.064, P = 0.001; SP-SP: Δ = −0.038, P = 0.024; IT-IT: Δ = −0.058, P = 0.001; all n = 19]. The same inversion was observed in another narrative (supplementary text S8).

**Fig 3.**
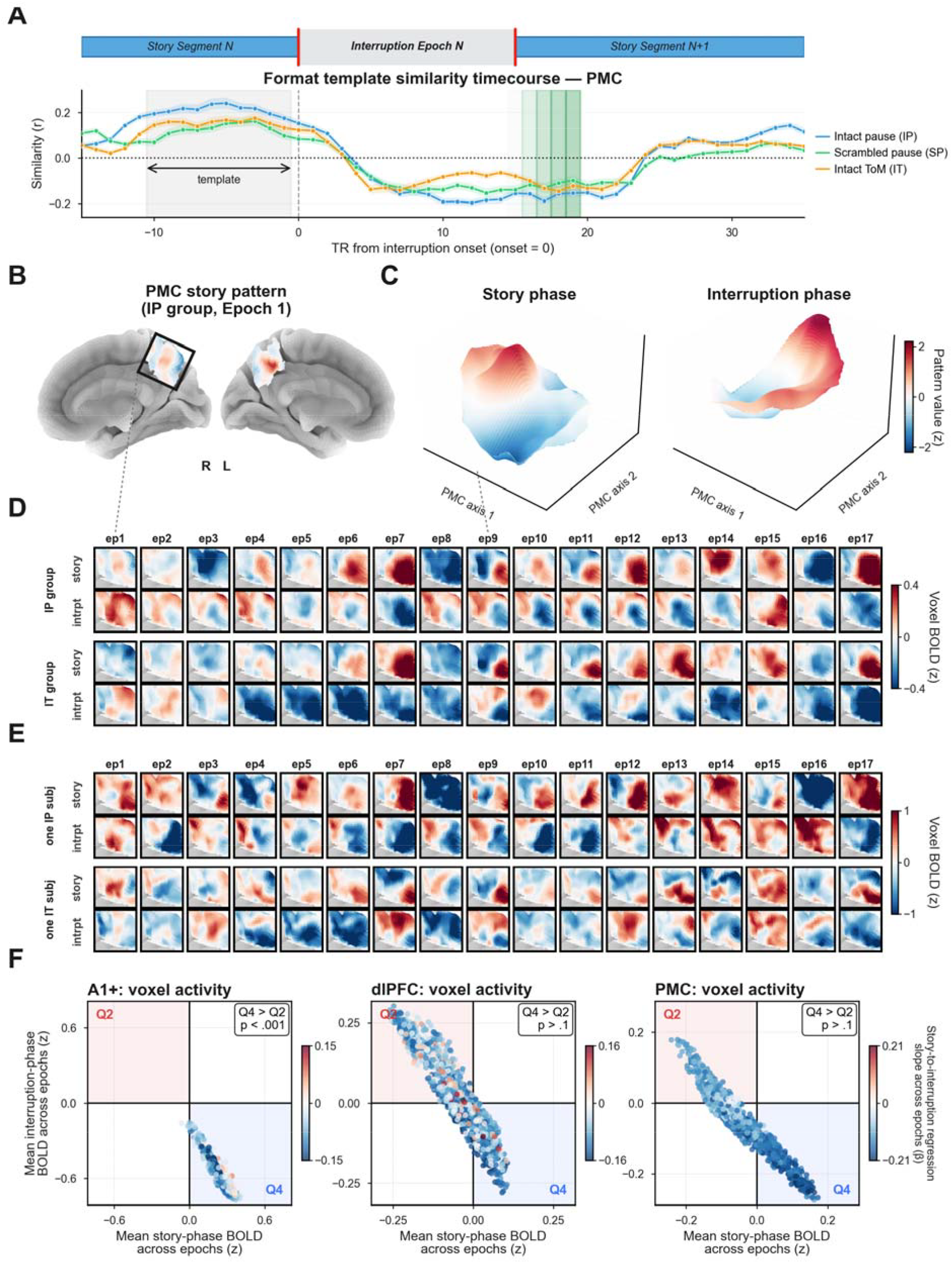
PMC sustains narrative context in a transformed format during interruption. (A) Pattern similarity between the preceding story-phase template and PMC patterns around interruption onset at each TR (mean ± SEM; n = 19 per condition): similarity turns negative during interruption and recovers after story resumption. Red/green shading marks time points contaminated by neighboring epochs or early story resumption; gray band, story-phase template window. (B) Group-mean storyphase PMC BOLD signal (IP group, Epoch 1) on the right (R) and left (L) medial surfaces; the boxed right-hemisphere region is expanded in the pattern maps below. (C) Group-mean PMC story- and interruption-phase patterns of Epoch 9 (IP group) as threedimensional topographies: height and color indicate each voxel’s pattern value, so peaks and troughs are directly visible. (D) Group-mean story- and interruption-phase patterns for all segments and epochs (IP and IT groups). (E) Example singleparticipant patterns (one IP, one IT). (F) Voxel-wise story-phase versus interruption-phase activity in A1+, dlPFC, and PMC, averaged across participants and epochs and pooled across interrupted conditions (n = 57). A post-stimulus undershoot predicts more voxels turning from positive to negative during interruption (Q4) than the reverse (Q2). Dot color shows each voxel’s storyto-interruption regression slope (red, sign-preserving; blue, sign-reversing).

#### PMC pattern transformation was robust to alternative explanations based on hemodynamic undershoot, normalization, and preprocessing

First, to test whether the effect reflected a poststimulus hemodynamic undershoot or neural adaptation (*21, 22*), we quantified the direction of voxel-wise sign changes from story to interruption. Unlike auditory and language regions where the pattern was more consistent with undershoot, PMC showed comparable positive-to-negative and negative-to-positive inversions, (Fig. 3; supplementary text S10). Second, because whole-run temporal z-scoring could in principle induce pattern anticorrelation between story and interruption phases, we confirmed that the story-to-interruption correlation remained negative after separately z-scoring the story and interruption phases (supplementary text S11). Finally, because temporal filtering can introduce anticorrelation in adjacent task phases (*23, 24*), we re-preprocessed the data without high-pass filtering or nuisance regression, separately z-scoring the two phases, and repeated the analysis. The signal reliability in the unfiltered data was lower, but we again observed epoch selective PMC patterns during the interruption, with a negative storyto-interruption correlation (supplementary text S12) with the shape of TR-by-TR lag-correlations closely preserved (supplementary text S13). Together, these controls indicate that PMC pattern transformation was not explained by normalization, hemodynamic undershoot, or preprocessing artifacts.

These findings suggest that PMC preserves interrupted narrative context in a transformed format that is detectable even during the processing of an alternate task.

### PMC pattern persistence predicted neural resumption and narrative memory

If the PMC background code supports mental continuity, its persistence during interruption should predict how readily participants resume the narrative. Therefore, we tested whether shared PMC pattern sustained across interruption predicted neural resumption and later memory. Because hippocampal boundary responses support event coding (*19*) and could be relevant to later resumption, we further asked whether PMC persistence explained variance beyond hippocampal activity at interruption onset. PMC persistence was quantified as within-condition ISPC across all TR pairs within matching interruption epochs (see supplementary text S16 for consistent results from alternative measures of persistence), and neural resumption was measured as each participant’s neural realignment with the continuous group in DMN regions (see materials and methods, Brain-behavior modeling).

#### Shared PMC persistence and hippocampal boundary activity both predicted neural resumption

In separate models (n = 57), stronger PMC persistence predicted greater DMN pattern realignment [b = 0.386, t(53) = 4.99, P < 0.001], as did greater hippocampal boundary activity [b = 0.093, t(53) = 2.91, P = 0.005]; both remained significant in the same model [PMC: b = 0.350, t(52) = 4.67, P < 0.001; hippocampus: b = 0.069, t(52) = 2.50, P = 0.016], suggesting partly independent contributions to neural resumption (Fig. 4). Moreover, PMC persistence during interruption predicted additional variance of neural resumption beyond what was predicted by preceding story-phase ISPC (supplementary text S17).

**Fig 4.**
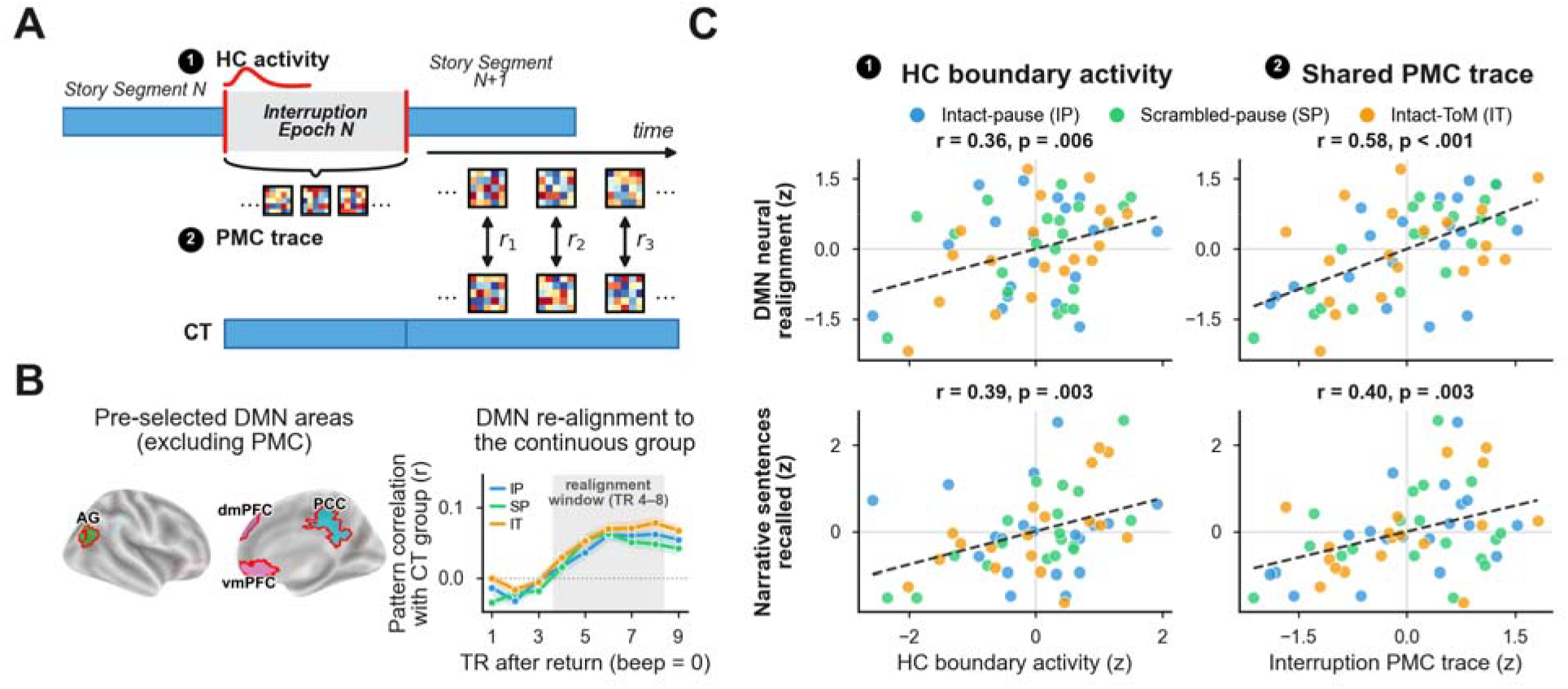
Hippocampal and cortical pathways to mental continuity. (A) Schematic of two candidate pathways for carrying context across interruption: hippocampal boundary activity at interruption onset (1) and a persistent PMC trace during interruption (2). After story resumes, neural resumption is measured as each participant’s default-network realignment with the continuous-listening group. (B) The default network ROIs used to measure neural realignment and the post-return realignment time course for each condition (mean ± SEM); the default network ROIs exclude the PMC ROI to avoid circularity; the realignment window spans TRs 4 to 8 after story resumption. (C) Hippocampal boundary activity and PMC pattern persistence in relation to neural resumption (top) and the proportion of narrative sentences recalled (bottom). Dots are participants (n = 57 for neural resumption; n = 55 for recall); values are z-scored within condition for visualization, so both axes are in standard-deviation (z) units and scores below the condition mean are negative. Statistical inference used pooled ordinary-least-squares models with condition entered as a fixed effect.

#### Shared PMC persistence and hippocampal boundary activity both predicted narrative memory

In separate models (n = 55), PMC persistence predicted recall [b = 0.764, t(51) = 3.00, P = 0.004], and hippocampal boundary activity also predicted recall when tested alone [b = 0.232, t(51) = 2.47, P = 0.017]. When both were entered together, both predictors remained significant [PMC: b = 0.663, t(50) = 2.63, P = 0.011; hippocampus: b = 0.185, t(50) = 2.04, P = 0.047].

Overall, these findings link greater shared PMC pattern persistence during interruption to more canonical post-interruption dynamics and more accurate narrative memory, and the PMC persistence effects complement the effects of hippocampal activity at interruption onset.

## Discussion

How do humans maintain mental continuity in the face of continual interruption? One pathway is by retrieving snapshots of the past: mental context can be stored via hippocampal-cortical interactions at the start of an interrupting event (*18, 19*) and then reinstated when the interruption is over. However, it has long been hypothesized that additional mechanisms sustain mental context through interruptions, enabling more rapid and automatic retrieval of the prior context (*4, 25*). We have identified a background process that maintains high-level narrative context in a rotated format within the posterior-medial cortex (PMC). This PMC representation, which changed gradually from stage to stage of the unfolding narrative, persisted in the face of concurrent tasks and predicted successful neural processing and behavioral recall of interrupted narratives.

Default network regions are functionally and anatomically situated to provide high-level mental continuity. These systems represent and integrate abstract episodic and narrative information over tens of seconds (*5–7, 10, 26*) and they are closely coupled to medial temporal lobe memory systems (*27, 28*). When people were interrupted during narrative processing, we observed a transient increase in both hippocampal activity (Figure 1H) and PMC activity (supplementary text S2); hippocampal activity (*18, 19*) and hippocampal-PMC coupling (*29*) are linked to superior event coding. Immediately following these transient increases, we observed the stable and reliable PMC multivoxel pattern which was selective to the preceding narrative epoch.

The PMC interruption patterns were consistent with a gradually evolving narrative context: epochs closer together in the narrative exhibited more similar interruption patterns, but only when the narrative was presented in its original order. Moreover, this gradual change did not simply reflect gradual shifts in the local semantic content of the story sentences preceding each interruption: we did not observe the gradual change when we re-ordered the scrambled-pause epochs into the intact narrative sequence (Fig. 2D).

The process we observed in PMC is different from conventional working memory in three respects. First, we found the neural signature within default-network systems, which are generally involved in internal, self-related and situational representations. Second, the PMC patterns retained information about the narrative even when active cognition was occupied elsewhere during a competing theory-of-mind task. This is inconsistent with a standard effortful-maintenance account and accords with evidence that narrative content can linger beyond external input and that high-level narrative integration can persist despite severe hippocampal damage (*2, 11*). Third, the stable pattern observed during interruption was a transformation of the preceding story-phase pattern. This phenomenon is distinct from standard reinstatement, in which prior content returns in a similar neural format, and from persistent activity, in which the relevant pattern remains online.

The negative correlation we observed can be understood as a rotation or inversion of the neural state. Such transformations have been observed in visual cortex when visual working memory traces are deprioritized (*14, 15, 30*) and when individual events are encoded as part of a complex sequence (*31*). The rotated fMRI pattern may reflect changes in the underlying neuronal populations contributing to the blood-oxygen-level-dependent (BOLD) signal within each voxel

(*32*) or a transition between coding subspaces to reduce interference (*13*). A recent EEG decoding study of visual working memory showed suppression of the decodable trace of no-longer-prioritized content, and concluded that it arose from a hijacked perceptual adaptation controlled by top-down modulation (*33*). Although precisely quantifying the transformation will likely require single-neuron recordings, these data establish that a background representation of narrative information persists through interruption in a format distinct from the story phase.

Hippocampal mechanisms may bind or index specific event content at moments of disruption, while PMC preserves the broader narrative scaffold across the intervening period. We found that hippocampal activity increased at interruption boundaries, and this increase independently predicted later neural resumption and memory, consistent with boundary-related encoding and event indexing (*5, 18, 19*). In parallel, the persistence of the PMC patterns also predicted both the neural resumption strength and the memory effects. A crucial open question, then, is whether the PMC state reflects cortical-hippocampal coordination, or whether the PMC transformation operates independently.

Altogether, our findings indicate that mental continuity depends not only on hippocampal encoding and retrieval, but also on a PMC background representation which survives interference. More broadly, such a background process may help explain how we can maintain one mental world while engaging with another. A person listening to an audiobook on the way to work, for example, can carry the unfolding narrative world in mind while navigating and acting within the physical world. By preserving high-level context in a transformed format while other processing takes priority, the default network may provide a mechanism for keeping an internally constructed world available alongside ongoing experience of the external environment.

## Acknowledgments

We thank Shima Rahimi Moghaddam for assistance with stimulus development and testing; Savannah Born, Arthur Li, Hanna Fu, and Rocio Mayorga for feedback on the stimuli and assistance with fMRI data collection; Janice Chen and Gabriel Kressin Palacios for assistance with fMRI data preprocessing. Claude (Anthropic; Claude Opus 4.8, Claude Opus 5, Claude Fable 5, and Claude Fable 5.1) assisted, under the authors’ direction, with organizing existing analysis code, rendering author-designed schematics, and editorial checks of author-written text; no generative-image model was used. AI tools were not used to formulate hypotheses, design the study or analyses, collect or analyze data, make inferential decisions, or interpret results; all scientific content was produced and reviewed by the authors (Supplementary Materials, Use of large language models).

## Author contributions

Conceptualization: X.L., C.J.H.

Methodology: X.L., H.L., C.J.H.

Software: X.L.

Formal analysis: X.L.

Investigation: X.L.

Data curation: X.L.

Visualization: X.L.

Project administration: X.L.

Writing – original draft: X.L.

Writing – review & editing: X.L., H.L., C.J.H.

Funding acquisition: C.J.H.

Supervision: C.J.H.

## Data, code, and materials availability

All code and analysis-ready data required to reproduce the analyses and figures reported in the manuscript and Supplementary Materials are publicly available. Analysis and figure-generation code is available at https://github.com/xianNeuro/mental_continuity. A frozen version of the code used for this study, together with the required input and the corresponding output is permanently archived on Zenodo (*34*). The preprocessed fMRI data will be deposited on OpenNeuro upon publication. Stimulus transcripts, interruption boundaries, interruption timings, scrambling manipulations, and theory-of-mind probe materials are included in the archived data.

## List of Supplementary Materials

Materials and Methods

Supplementary Text

Figs. S1 to S13

Tables S1 to S4

References (35-47)

