## supplementary materials for "Cortical and Hippocampal Pathways to Mental Continuity"

**This PDF file includes:**

**Materials and Methods**

**Supplementary Text**

Figs. S1 to S13

Tables S1 to S4

Other Supplementary Material for this manuscript includes the following:

MDAR Reproducibility Checklist

**Materials and Methods**

***Participants***

Ninety-one right-handed native-English-speaking adults (33 male, 58 female; ages 18-40 years, mean 23.3 years) with normal or corrected-to-normal vision and hearing participated in the fMRI experiment. Eighteen of the 91 participants were excluded from further analysis in this study due to data quality issues: in-scanner sleep during task (n = 5); head motion persistently exceeding 3 mm absolute displacement (*n* = 2); extremely short free recall (audio length < 3 min) indicating inattentiveness during encoding (*n* = 7); incomplete scan due to technical issues (*n* = 1); participant requested to exit the study (*n* = 1); neural data failed quality control check requiring high inter-subject synchrony in auditory processing areas (n = 1); low recognition memory (four-alternative forced-choice accuracy below chance) (n = 1). These exclusion criteria were fixed during the initial data-quality screening of the dataset, before any of the pattern or brain-behavior analyses were performed. A further four participants ended the scanning session early, at their own request or because of a technical interruption, and one pilot participant was scanned with earlier stimulus timing; none of these five yielded a usable dataset and they are not included in the 91. This yielded a final sample of 73 participants (28 male, 45 female, mean age 22.8 years, SD 4.1, range 18-40; *n* = 16 for the CT condition; *n* = 19 for each of the IP, IT and SP conditions).

All procedures were approved by the Johns Hopkins Medicine Institutional Review Boards, and all participants provided written informed consent. Group size was designed in advance to 15-20 participants based on previous naturalistic fMRI studies (5, 6). Participants were compensated $30 per hour for scanning and $15 per hour for the behavioral session.

***Conditions***

Each participant listened to two narratives, and each narrative could be drawn from one of four conditions: continuous (CT), intact order with pauses (IP), intact order with theory-of-mind interruptions (IT), and scrambled order with pauses (SP). In the CT condition, participants listened to the narrative without any interruptions. This served as the reference condition, establishing the typical neural patterns at each time point when each part of the stimulus was presented without interruption and with appropriate narrative context. In the IP condition, participants listened to the narrative in intact sequence but interrupted by silent pauses that lasted for 22.5-28.5 seconds. The IP condition provided interruption phases in which we could search for neural activity locked to the preceding narrative context, but without any immediate stimulus drive. In the IT condition, participants listened to the narrative in intact sequence, just as in the IP condition and with the same number and duration of interruptions, but each interruption phase was filled by a story-irrelevant auditory vignette that described a scenario, followed by a yes/no question that required theory-of-mind type (ToM) situational reasoning based on the vignette’s content. This condition was designed to interfere with participants’ working memory for the main narrative by requiring them to operate on a story-irrelevant situation. In the SP condition, participants listened to the narrative in a scrambled sequence, with the same number and duration of silent pause interruptions as the IP group. The scrambling of the narrative was designed to disrupt the formation of a coherent narrative context, while preserving local semantic and situational content on the scale of tens of seconds.

***Stimuli***

The main narrative was a murder mystery titled "So Much Water So Close to Home" by Raymond Carver, professionally narrated by a voice actress, lasting 14 minutes and 9 seconds (2,354 words). A secondary narrative was a 9 min 9 seconds audio of stand-up-style live-storytelling monologue (1,429 words), titled "Not the Fall That Gets You" by Andy Christie.

***Narrative segments.*** In order to determine the location of the interruptions, narratives were segmented into units of 30-60 seconds. This rendered 18 segments in the main narrative (“So Much Water So Close to Home”), each containing a mean of 13.3 sentences (range 8-20) and 130 words (range 105-165). The same process applied to the secondary narrative (“Not the Fall That Gets You”) yielded 12 segments, each containing a mean of 9.6 sentences (range 5-13) and 120 words (range 77-164). The segment boundaries were designed to fall at natural sentence boundaries never interrupting the middle of a sentence, but with the primary consideration being the audio duration and word counts in the story segments. Thus, these segment boundaries were not chosen to match event boundaries, although some boundaries may overlap with event boundaries in the original narrative. In the three interruption conditions (IP, IT, SP), these story segments rendered 17 locations to insert the interruption epochs in the main narrative, and 11 in the secondary narrative. Across all interruption conditions, the transition from the story phase to the interruption phase was cued by a clear double-beep sound, and the same audio cue was applied at return to the story.

***Scrambled narrative (SP condition).*** For the scrambled condition, each narrative segment was further sub-divided into 3 finer sub-sections (“head”, “middle”, “tail”; segment 7 into four, giving 55 in total) at natural sentence boundaries. The division was primarily based on length (a mean of 4.4 sentences per sub-section, range 1-10; mean of 43 words per sub-section, range 21-92). The sub-sections were reordered to make the story difficult to follow, while preserving the sub-sections that immediately preceded and followed each segment boundary (Fig. S1). The reordering was constructed by hand under an explicit constraint: wherever possible, sub-sections adjacent in the scrambled run came from non-adjacent original segments and from different within-segment positions, so that consecutive sub-sections could not be readily integrated against one another when heard in the scrambled condition. At the same time, the sub-section pair flanking each interruption (i.e., the tail of segment N and the head of segment N+1) were kept adjacent: in this way, the local content immediately before and after every interruption was identical to that in the CT, IP, and IT conditions (Fig. S1). Overall, this procedure was designed to disrupt the global narrative coherence while preserving the per-sub-segment local audio content and local semantic comprehension to match that of the intact narratives.

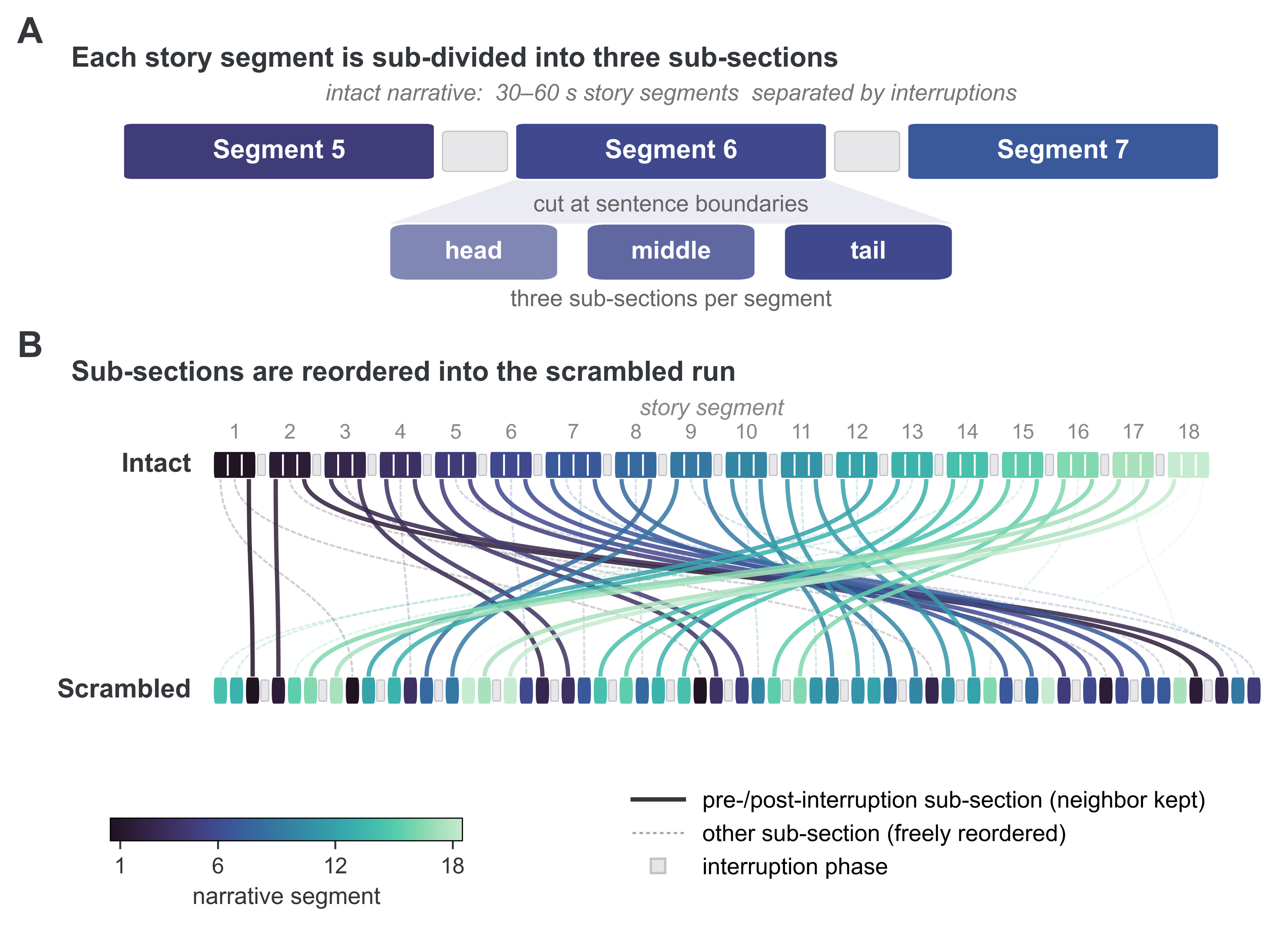

**Fig. S1.** Procedure for scrambling sub-sections to generate the stimulus for the scrambled-pause (SP) variant of the “So Much Water So Close to Home” narrative. Story segments are shown in blue-green shades; interruption phases are indicated by gray blocks. (A) Each 30-60 s story segment was cut at sentence boundaries into a head, middle and tail sub-section; Segment 6 is shown with its neighbors. (B) The full permutation of the 18 segments and 55 sub-sections is shown as a mapping from the intact-pause stimulus (top) to the scrambled-pause stimulus (bottom). Bold solid ribbons indicate the tails and heads of segments (which are shuffled alongside their intervening interruption phase); faint dotted ribbons track the middle sub-sections which could be shuffled to any other middle position.

***Theory-of-mind probes (IT condition).*** The interruption epochs of the IT condition were filled with short, narrative-irrelevant situational reasoning vignettes adapted from the Saxe Lab False-Belief Localizer (*35*). Each probe consisted of a brief descriptive vignette introducing a character and a situation. The vignettes started immediately following the beep cue that marked the onset of the interruption and were followed by a single yes/no comprehension question about the character's belief (12 questions) or the veridical state of the scenario (5 questions). Participants responded by button press during a 4.5 s response window after the question and before the end-of-interruption beep cue. Seventeen distinct vignettes were used in the main narrative and 11 in the second narrative; all 11 second-narrative items were a subset of the main-narrative items. Consequently, to avoid repeated exposure to the same ToM questions, participants that were assigned to the IT condition in the main narrative were assigned to the IP condition for the second narrative. The vignettes and questions were lightly edited from the original localizer task for this study in two ways. First, we edited the questions to render a balance of Yes/No correct responses. Second, we shortened the vignettes and questions slightly so that each fit within the interruption epoch. The vignettes are known to engage activity within default mode network regions, and participants’ focus on and comprehension of the vignette were required to be able to answer the question correctly (Table S1).

Before scanning, participants were explicitly informed that they would listen to an audio story and that the story would be interrupted occasionally by a question regarding a separate situation, unrelated to the story. Participants were told that they should listen to the unrelated situation description and answer the question, and that the story would resume afterwards. Participants were informed that their memory for the story would be later tested, and that they should also try their best to answer the separate story-irrelevant questions correctly. Participants who received theory-of-mind interruptions during the main narrative answered its 17 interruption questions well above chance (mean acc = 0.916, SE = 0.020, range = [11/17, 17/17], n = 19), as did the participants who received the 11 interruption questions during the second narrative (mean acc = 0.884, SE = 0.021, range = [8/11, 11/11], n = 18). These results confirm that participants indeed directed their attention to the vignettes, which were unrelated to the main or secondary story.

**Table S1.** The 17 situational vignettes and questions used in the intact-ToM condition. For each vignette and question, we note the yes/no answer key (Y/N) and the narratives in which each was used. The 11 probes marked “Both narratives” were also used in the second narrative for participants who did not hear ToM questions in their first narrative; “Main only” probes were presented in the main narrative alone.

| Item | Vignette and question | Key | Used |
| --- | --- | --- | --- |
| 1 | The morning of the high school dance Sarah placed her high heel shoes under her dress and then went shopping. That afternoon, her sister borrowed the shoes and later put them under Sarah's bed. When Sarah gets ready, does she assume her shoes are under her dress? | Y | Both |
| 2 | John told Mary that he had lost his keys. The two of them searched the house with no luck. Then Mary went outside to look in the car. Suddenly John noticed his keys behind the sofa. By the time Mary comes in, does John know where his keys are? | Y | Both |
| 3 | Expecting the game to be postponed because of the rain, the Garcia family took the subway home. The score was tied, 3-3. During their commute the rain stopped, and the game soon ended with a score of 5-3. When the Garcia family arrives home, do they believe the score is 5-3? | N | Both |
| 4 | When Lisa left Jacob, he was deep asleep on the beach. A few minutes later a wave woke him. Seeing Lisa was gone, Jacob decided to go swimming. Does Lisa now believe that Jacob is asleep? | Y | Both |
| 5 | Larry chose a debated topic for his class paper due on Friday. The news on Thursday indicated that the debate had been solved, but Larry never read it. When Larry writes his paper, does he think the debate has been solved? | N | Both |
| 6 | A window wiper was commissioned by a CEO to wipe an entire building. He finished the right side, but his platform broke before he could do the left side. The next morning the CEO arrived with foreign investors. When the CEO comes to work, do they discover that all of the windows are cleaned? | N | Main |
| 7 | Sally and Greg called ahead of time to make a reservation for the back-country cabin. The park ranger forgot to write down the reservation and two other hikers got to the cabin first. When the hikers arrive, do they find their cabin unoccupied? | N | Both |
| 8 | Amy walked to work today. When George woke up, he saw her car in the drive. Her room was quiet and dark. George knows that when Amy is sick, she lies down in a dark room. Does George think Amy is sick today? | Y | Both |
| 9 | At night a bear broke into a cooler near a tent and drank the soda. Five hours later, the campers woke up and went to their cooler for breakfast. Do the campers find the cooler empty of soda? | Y | Both |
| 10 | Anne made lasagna in the blue dish. After Anne left, Ian came home and ate the lasagna. Then he filled the blue dish with spaghetti and replaced it in the fridge. Does Anne think the blue dish contains spaghetti? | N | Main |
| 11 | Jenny put her chocolate away in the cupboard. Then she went outside. Alan moved the chocolate from the cupboard into the fridge. Half an hour later, Jenny came back inside. Does Jenny expect to find her chocolate in the cupboard? | Y | Main |
| 12 | When the class' science test was handed back, Shannon was mistakenly given Adam's test. A large B was written on the front of Adam's test, but Shannon's actual grade was an A. In reality, did Shannon receive an A on the exam? | Y | Main |
| 13 | Every day Jill goes to the coffee shop on the corner and orders a latte, her favorite drink. Today, the cashier misunderstands Jill and prepares a mocha instead. Does Jill think her drink will taste like a mocha? | N | Both |
| 14 | Hopeful to catch a prize fish, George went fishing. That afternoon, he saw his fishing line bend over as if he had caught a big fish. Actually, George’s fishing pole had snagged a small tire. At the end of the fishing line, does George see a fish? | N | Main |
| 15 | The girls left ice cream in the freezer before they went to sleep. Over night the power to the kitchen was cut and the ice cream melted. When they get up, do the girls believe the ice cream is melted? | N | Main |
| 16 | Ken told Andrea that he was going shopping for sandals. At the shoe store, Ken noticed a very nice pair of boots on sale and bought them instead. Does Ken's shoe store bag contain sandals? | N | Both |
| 17 | When Jeff got ready this morning, he put on a light pink shirt instead of a white one. Jeff is colorblind, so he can't tell the difference between subtle shades of color. In reality, is Jeff's shirt pink? | Y | Both |

***Interruption-epoch timing.*** Interruption epochs in IP, IT, and SP were carefully designed to match in total number (17 for the main narrative, 11 for the second narrative) and the per-epoch duration. Each interruption epoch had the same duration across the three interruption conditions (IP, IT, and SP). The interruptions were matched across conditions so that they had identical preceding and following (tail and head) narrative content, rather than by serial position. This enabled a stimulus-matched and duration-matched comparison of neural dynamics in the SP condition against other conditions. The onset and offset of every interruption epoch were aligned to volume-acquisition (TR) boundaries (TR = 1.5 s), so that each interruption began and ended at the start of a TR and lasted an integer number of TRs. The Continuous condition contained no interruptions, but equivalent time windows in the intact narrative (defined by the same segment-boundary positions) were aligned to the IP, IT, and SP epochs for cross-condition analyses. The onset and offset of every interruption epoch were marked by a brief double-beep auditory cue: a 400 Hz beep for 150 ms, then 80 ms of silence, a second 400 Hz beep for 150 ms, then a further 1120 ms of silence. Thus, the onset/offset cues for each interruption were 1500 ms long, matching the TR of the BOLD acquisition, and ensuring that the following audio always began on a TR boundary.

***Experimental procedure***

Each participant heard two narratives. CT condition participants heard both narratives in the same continuous format with segments in their original order. SP participants heard both narratives in scrambled sequence. IP and IT participants were rotated across conditions for the two narratives: if a participant heard the IP version of the main narrative, they would hear the IT version of the second narrative stimulus, and vice versa. Within a single scanning session, each participant listened to the main narrative (“So Much Water so Close to Home”) first in one continuous fMRI run. They later heard the second narrative (“Not the Fall That Gets You”) in another continuous fMRI run. Finally, they recalled the main narrative aloud in a separate continuous free-recall run. A 120-second free-association task (*2*), in which participants spoke free word associations aloud to approximate spontaneous thought, was completed immediately before and after the main narrative and immediately after the second narrative, within the same continuous fMRI scan that contained each narrative. This allowed us to detect participant’s thoughts pre- and post-story listening, and to detect potential data quality issue such as falling asleep or inattentiveness. Immediately after the first and second narrative fMRI acquisitions, participants reported their immersion in the story (transportation) and the degree to which story-related thoughts had persisted in their minds. Each main-narrative run comprised 736 TRs in the CT condition and 1026 TRs in each of the interrupted conditions. Each second-narrative run comprised 456 TRs (CT) or 636 TRs (interrupted). All run durations included the free-association periods.

Before scanning, all participants were encouraged to stay attentive and were told that their memory for both narratives would be tested. Participants in the interrupted conditions received the same instructions as the CT condition, with additional instructions about the interruptions they would encounter: IP participants were told that silent, double-beep-marked pauses would punctuate the narrative; SP participants received the same instruction and were additionally warned that the narrative might be hard to follow but that they should stay attentive; IT participants were told that each interruption would pose a narrative-unrelated yes/no theory-of-mind question, and each answered one practice ToM probe (not used in the experiment) before scanning. Behavioral responses occurred only during the interruptions in the intact-ToM condition.

After scanning, and outside of the scanner room, participants completed a behavioral battery. Memory for each narrative was tested in two formats that targeted the story content immediately before and after the interruption epochs: a fill-in-the-blank (FIB) test of recollection, in which spans of two to eighteen words were removed from sentences for the participant to complete, and a four-alternative forced-choice (4AFC) test of recognition, in which a word or phrase was removed and four options were offered, one correct. Participants then completed a questionnaire on their overall comprehension of the two stories, followed by self-report questionnaires about their cognitive experience before, during, and after the interruptions, including how much they thought about the narrative during the interruption epochs. Recall and comprehension scores were compared across the four conditions with one-way analyses of variance, followed by pairwise Welch two-sample t-tests (unequal variances) with Bonferroni correction, run when the omnibus test reached P < 0.1.

***Behavioral data processing***

***Human ratings for naturalistic stimuli.*** Our laboratory maintained a team of trained human raters who routinely performed processing and rating tasks on naturalistic stimuli, including speech transcription and recall scoring. The raters were trained and supervised by a graduate-student team leader, who reviewed their work for quality and provided ongoing feedback. All raters followed a standardized written instruction set to ensure consistency across coders.

***Transcription and segmentation of free recall.*** The audio recording of the free-recall for the main narrative was transcribed automatically using Whisper (Large-v2 model; OpenAI), with the automatic transcripts subsequently manually corrected by trained raters. Each transcript was segmented into sentences (or parts of sentences), and timestamps were identified for the beginning and end of each segment. Transcribed segments that ended before the beginning of the recall run, or that began after its end, were excluded from analysis. Filler words and disfluencies (e.g., “um”, “uh”, “ah”) were removed during transcription, so that the resulting text reflected the narrative content the participant produced rather than speech-production artifacts.

***Matching recall to story sentences.*** Prior to the experiment, the text of the main narrative was divided into 240 sentences. Participant recalls were segmented based on both punctuation and topic shifts, corresponding to sentences or parts of sentences. Each recall segment was then assigned to the story sentence or sentences it referred to, with a recall segment potentially aligning with none, one, or multiple story sentences. A story sentence was scored as recalled if at least one recall segment referred to it (binary 0/1 per sentence). The per-participant recall score used in the brain-behavior analyses was the proportion of the 240 main-narrative sentences recalled.

***fMRI data collection and preprocess***

***MRI data acquisition.*** MRI scanning was conducted at the F. M. Kirby Research Center for Functional Brain Imaging at Kennedy Krieger Institute on a 3 Tesla Philips Ingenia Elition scanner with a 32-channel head coil. Functional images were acquired using a T2*-weighted multiband accelerated echo-planar imaging (EPI) sequence (TR = 1.5 s; TE = 30 ms; flip angle = 52°; acceleration factor = 4; 60 oblique axial slices; grid size 112 × 112; voxel size 2 × 2 × 2 mm^3^). Fieldmap images were also acquired to correct for B0 magnetic field inhomogeneity (60 oblique axial slices; grid size 112 × 112; voxel size 2 × 2 × 2 mm^3^). Whole-brain high-resolution anatomical images were acquired using a T1-weighted MPRAGE pulse sequence (150 axial slices; grid size 224 × 224; voxel size 1 × 1 × 1 mm^3^).

***fMRI preprocessing (primary)***. Preprocessing was performed in FSL 6.0 (*36*), including motion correction, high-pass filtering (140 s cutoff), and coregistration and affine transformation of the functional volumes to a template brain (Montreal Neurological Institute (MNI) standard). Functional images were spatially smoothed with a Gaussian kernel of 4 mm full width at half maximum (FWHM) in 2 mm template space and then resampled to 3 mm isotropic voxels for all analyses. All calculations were performed in volume space. Projections onto a cortical surface were performed for visualization only.

***fMRI preprocessing (secondary)***. In order to rule out the possibility that interruption phase patterns reflect preprocessing artifacts, we re-processed the data with a separate minimal pipeline and conducted a series of control analyses. For the control analyses of the PMC story-to-interruption transformation (Sections S12 and S13), functional images were re-preprocessed using fMRIPrep (version 21.0.2; *37*) with default settings. Functional images were corrected for head motion and B0 magnetic inhomogeneity using the acquired field maps, co-registered using rigid-body, six-degree-of-freedom transformation to the anatomical image and transformed to the MNI152NLin6Asym template. The resampling to 3 mm isotropic voxels and spatial smoothing at 4 mm FWHM was performed at the multivoxel-pattern extraction stage to match the main FSL pipeline. No high-pass filtering or nuisance regression was applied, so that the transformation could be evaluated on minimally processed data.

***Regions of interest (ROIs)***

We pre-defined a collection of ROIs based on our research goal and prior literature. The pre-selection covered three functional families: default-mode subregions that were the primary substrate of the predicted “background” representation of episodic content; working-memory areas that tested a more conventional active-maintenance account; and long-term-memory regions that tested the boundary-encoding account. Early auditory and language pathway regions were also included as control regions that should primarily track local properties of the auditory narrative. Each ROI mask was created by combining relevant parcels from the Schaefer 400-parcel, 17-network atlas (*38*) to maximally match with the time-course analysis extracted from the whole-brain (Section S1). The bilateral hippocampal ROI was defined from the Tian subcortical atlas (39) by combining its anterior and posterior hippocampal subdivisions into a single mask.

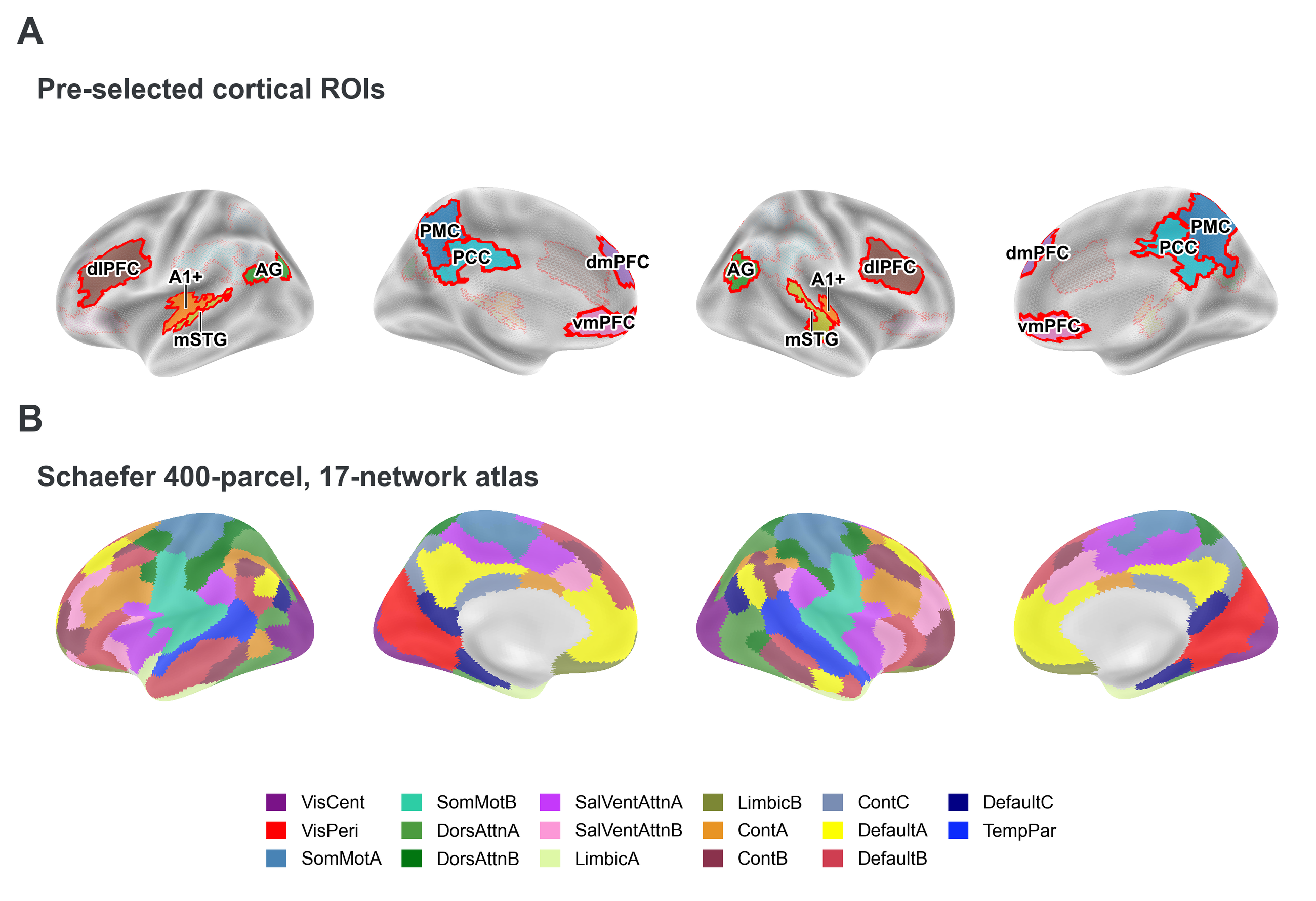

**Fig. S2.** Regions of interest and the whole-brain parcellation, on the fsaverage inflated surface. (A) The eight pre-selected cortical ROIs: posterior medial cortex (PMC), posterior cingulate cortex (PCC), angular gyrus (AG), dorsomedial prefrontal cortex (dmPFC), ventromedial prefrontal cortex (vmPFC), early auditory cortex (A1+), middle superior temporal gyrus (mSTG), and dorsolateral prefrontal cortex (dlPFC). Each ROI is shown as a colored fill outlined in red. The hippocampal ROI is not shown in this surface representation. (B) The Schaefer 400-parcel, 17-network atlas (*38*) used for the whole-brain analyses, rendered from the official FreeSurfer surface annotation and colored according to Yeo 17-network membership (see legend).

***Default-mode sub-regions.*** Five cortical sub-regions of the default-mode system, each generated by taking the union of parcels from the Schaefer 400-parcel atlas, were selected to test the prediction of a “background” representation within DMN regions: posterior medial cortex (PMC; dorsal precuneus and adjacent posterior cingulate, which also draws in neighboring control- and salience-network precuneus parcels), posterior cingulate cortex (PCC), angular gyrus (AG), dorsomedial prefrontal cortex (dmPFC), and ventromedial prefrontal cortex (vmPFC). These regions are implicated in long-timescale narrative integration and representation of event-structure and narrative content (5, 6, 10, 40). We predicted that, within these regions, we would identify a subset whose representations were shared across participants, epoch-selective, evolving across coherent narrative time, and preserved when attention was diverted to the unrelated ToM task.

***Working-memory and long-term-memory regions.*** Dorsolateral prefrontal cortex (dlPFC) was included as candidate substrates for a conventional active-maintenance account (41) as well as a candidate site for an “episodic buffer” (42); it is thought to be involved in working memory and effortful control over completing tasks (*43*). The hippocampus (HC) is necessary to encode new long-term memories, and shows large transient activity increases at major event offsets (18, 19, 44). Therefore, we included a hippocampal ROI to quantify how much hippocampal-cortical boundary encoding at the onset of interruption predicts later retrieval and resumption processes. We predicted a transient hippocampal BOLD increase at interruption onsets, consistent with prior literature, but distinct from the sustained background pattern which we expected in cortical default network regions.

***Auditory and language pathway regions.*** Early auditory cortex (A1+) and middle superior temporal gyrus (mSTG) were included as positive controls for stimulus-driven processing during story listening (7, 45). In these regions we expected reliable and epoch-selective neural pattern correlations during the story phase, but not during the interruption phase.

***Analysis methods***

***fMRI data extraction and processing.*** All neural analyses operated on multivoxel activity patterns taken from the preprocessed functional time series in 3 mm MNI152 space (see fMRI preprocessing). For each voxel within the mask of each ROI, we had a single time course of BOLD values, one per TR, from each continuous scan. Before any pattern correlations were computed, each voxel’s time course was temporally z-scored across all TRs of the run. This whole-run, per-voxel z-scoring placed every voxel on a common scale, so that the inter-subject pattern correlations reflected the spatial patterning of activity rather than voxel-wise differences in the temporal mean or variance of BOLD signal. This whole-scan z-scoring was used throughout the manuscript, except for the control analyses that specifically tested the influence of this procedure. The supplemental analyses that separately standardized the story and interruption phases are described alongside the pattern-transformation controls in Sections S11 and S12.

***Whole-brain analyses.*** As supplemental analyses providing a whole-brain view beyond the preselected ROIs, each of the main-text pattern analyses was repeated across all parcels of the Schaefer 400-parcel, 17-network atlas (38). For the whole-brain time-course analyses (Supplementary texts S1 and S2), each voxel’s time course was z-scored across the run and then averaged within each parcel. Multivoxel-pattern analyses used the same whole-run voxel-wise z-scoring within each parcel as the corresponding ROI analyses (Sections S6 and S15). These analyses yielded whole-brain maps of the effects reported in the predefined ROIs. Results are reported with false-discovery-rate correction across the 400 parcels. Where stated, we also generated some plots at a more lenient uncorrected threshold to provide a broader picture of the spatial distribution of graded effects.

***Hippocampal boundary activity.*** For each interruption epoch and participant, the HC boundary response was defined as the mean BOLD activity across the 5 TRs following the onset TR minus the mean activity across the 5 TRs immediately preceding onset. Boundary responses were then averaged across epochs to obtain one value per participant. To test for an overall hippocampal boundary response at interruption onset, boundary responses from all interrupted participants (IP, SP, and IT; n = 57) were tested against zero using a one-sample t-test.

***Time-time inter-subject pattern correlation.*** All pattern analyses were derived from a common time-time inter-subject pattern correlation (TTC) map (5, 6). For each region, the multivoxel pattern of one participant at each TR was correlated with the mean pattern of all other participants in the group at every TR, yielding a time-by-time matrix of inter-subject pattern similarity. A submatrix of time-time correlation patterns was then extracted around each interruption onset (one for each epoch of the narrative, with t = 0 locked to the onset of the interruption), and these time-time submatrices were averaged across epochs to generate the summary view (Figure 2A). For matching epochs, diagonal cells quantify pattern similarity across participants at corresponding time points, whereas off-diagonal cells quantify similarity across different time points and therefore indicate whether shared patterns persist or change over time. Mean TTC maps for matching and mismatching epochs under the different inter-subject correlation schemes are shown in Figure 2B. The analyses below quantify specific windows and comparisons within these maps.

***Pattern reliability analysis.*** We tested whether the multivoxel patterns during interruption epochs were reliably shared across participants. Inter-subject pattern correlation (ISPC) was measured in a 15-s window (10 TRs) starting 5 TRs from the interruption onset. The first 5 TRs (including the onset TR) of each epoch were excluded to minimize the immediate hemodynamic carry-over from the preceding story segment. For each participant and epoch, the participant’s multivoxel pattern was correlated at each TR with the mean pattern of the other participants in the comparison group. Correlations were averaged across the window to derive one participant-by-epoch ISPC value and Fisher-z transformed before averaging. We evaluated three within-condition comparisons (IP-IP; SP-SP; IT-IT) and one across-condition comparison (IT-IP), in which each IT participant was compared with the mean pattern of the IP group.

The resulting participant-level ISPC values were mutually dependent because each participant’s ISPC value was computed relative to the group-average pattern from the rest. We therefore assessed reliability using a delete-one-subject jackknife followed by a sign-flip permutation test, with participants as the unit of resampling (46). For each participant, *j*, we computed the jackknife pseudo-value

$\tilde{\theta}_{j} = n \cdot\hat{\theta} - (n - 1) \cdot\hat{\theta}_{-j}$,

where n is the number of participants, $\hat{\theta}$ is the full group reliability estimate (mean Fisher-z ISPC across participants and epochs), and $\hat{\theta}_{-j}$ is the corresponding estimate recomputed after removing participant j entirely. (thus, recomputing every remaining participant’s ISPC values). The resulting pseudo-values provided participant-level estimates of the contribution to the group statistic while accounting for the dependence introduced by the shared group-average pattern. To test whether reliability was greater than zero, we then sign-flipped the pseudo-values across participants over 10,000 permutations and recorded their mean on each iteration. The one-sided P value was calculated as (k + 1)/(N + 1), where k is the number of permuted means greater than or equal to the observed group mean and N = 10,000 (46).

***Pattern selectivity analysis.*** For each ROI we tested whether interruption patterns were specific to epochs, i.e., epoch selective. If the interruption patterns were epoch-selective, then the matching epoch pairs’ ISPC should be greater than the mismatching epoch pairs. Inter-subject pattern correlation (ISPC) was measured for the 10 TRs that began 5 TRs from interruption onset between matching and mismatching epoch pairs. For each participant and each epoch, the Pearson correlation between that participant's pattern and the mean pattern from all other participants in that group was computed at every TR in the epoch window and averaged across the 10 TRs to a single per-participant per-epoch score; selectivity statistics operated on these raw correlation values. For participant j, selectivity was defined as

Δ_j_ = r_match,j_ − r_mismatch,j_ ,

where r_match,j_  is the mean ISPC for matching epoch pairs and r_mismatch,j_ is the mean ISPC across mismatching epoch pairs. Group selectivity was defined as the mean Δ_j_ across participants.

We computed two additional across-condition comparisons of each SP participant to the IP group mean: first, we aligned the conditions’ serial position in which events were presented to participants during the experiment (SP-IP); second, we re-ordered the SP epochs to their original positions in the intact narrative before comparison (SP-IP-unscrambled). In the second variation, the analysis ensured that the interruption epochs being compared were preceded by the same “tail” section of story content (Fig. S1).

Because ISPC values derived from a shared group-average pattern are statistically dependent, significance was assessed using an epoch-label permutation test on the group-mean selectivity statistic (46). Under the null hypothesis that interruption patterns contain no epoch-specific information, the matching and mismatching labels are exchangeable. Within each participant, the matching and mismatching ISPC values were pooled and randomly reassigned to sets of the original sizes 10,000 times, and a “matching minus mismatching” selectivity statistic was recomputed after each permutation. Group-mean selectivity was recorded on each iteration to form a null distribution. The observed selectivity was evaluated using the one-sided permutation test predicting greater ISPC for matching than mismatching epochs, with P = (k + 1)/(N + 1), where k is the number of permuted values greater than or equal to the observed value and N = 10,000. This procedure preserves the within-participant data structure and tests whether interruption patterns shared across participants carry epoch-specific information. Between-condition differences were assessed using two-sided Welch two-sample t-tests on participant-level selectivity scores, with Welch-Satterthwaite degrees of freedom, because IP and SP data comprised independent groups of participants.

***Pattern evolution analysis.*** For each ROI, we tested whether the shared interruption pattern evolved over the course of the narrative. If the shared pattern tracked the unfolding narrative, inter-subject pattern correlation (ISPC) between two interruption epochs should decline as the distance between them increased. ISPC was measured for the 10 TRs beginning 5 TRs from interruption onset for every forward pair of interruption epochs. For each participant and each epoch pair, the participant's multivoxel pattern at the earlier epoch was correlated at each TR with the mean pattern of the other participants in the comparison condition (either within the same condition or another group) at the later epoch. Correlations were averaged across the 10 TRs to obtain a single participant-by-epoch-pair score; pattern evolution analyses were performed on these raw correlation values.

For each participant *j*, pattern-evolution was quantified as the slope *b*_j_ from an ordinary-least-squares regression, across epoch distances, of that participant’s mean ISPC at each distance on epoch distance. Epoch distances were restricted to approximately half the number of total epochs in each narrative, 1-8 for the main narrative (17 epochs) and 1-5 for the secondary narrative (11 epochs). Within these ranges, every epoch contributed to each modeled distance. At greater distances, epochs near the center of the narrative were progressively excluded and the estimates were based on increasingly fewer epoch pairs, yielding less representative estimates of narrative-wide pattern evolution (47). Distance of 0 corresponds to matching epoch pairs and was excluded from the slope estimate, so that the slope reflects the gradual change across epoch distances (all distances are mismatched) rather than a mean difference between matching and mismatching epochs.

Because the ISPC values are statistically dependent, slope significance was assessed using a within-participant epoch-distance-label permutation test on the group-mean slope (46). Under the null hypothesis that epoch distance is unrelated to the pattern similarity, epoch-distance labels are exchangeable within each participant. For each of 10,000 permutations, distance labels were randomly shuffled across each participant’s per-distance mean ISPC values, participant-level slopes were refit, and the resulting group-mean slope was recorded to form the null distribution. The observed slope was evaluated using a two-sided permutation test, with P = (k + 1)/(N + 1), where k is the number of permuted group-mean slopes whose absolute value was greater than or equal to that of the observed slope and N = 10,000. This procedure preserved the within-participant dependence structure of the interruption patterns across epochs while examining whether the shared interruption pattern changes progressively across the narrative. A negative slope indicates decreasing pattern similarity with increasing epoch distance. For the scrambled-pause condition the same model was additionally fit after reordering the epochs according to their positions in the intact narrative (SP-SP-unscrambled), such that epoch distance reflected the distance in the intact narrative sequence rather than the scrambled presentation order.

***Sequential testing procedure.*** The three pattern tests were applied in a fixed sequence. Pattern selectivity was evaluated only in regions with reliable interruption patterns (sign-flip P < 0.05), and pattern evolution was evaluated only in regions that were both reliable and epoch-selective. Cells not reached under this procedure are reported as not tested rather than as null results. For the whole-brain maps, a parcel was counted as showing the full profile when it passed all three tests at an uncorrected P < 0.005. The sequential testing procedure was used for the main-narrative ROI-screening analyses. For the second-narrative replication in the already identified PMC ROI (Sections S7, S8), all three predefined tests were repeated regardless of whether the preceding test reached significance, because the purpose was to evaluate replication of the full main-narrative profile rather than to select regions.

***Story-to-interruption transformation analysis.*** For ROIs showing reliable and epoch-selective interruption patterns, we tested the relationship between the interruption pattern and the pattern in the preceding story phase. For each participant and epoch, a story template pattern was computed by averaging across the 10 TRs immediately before interruption onset. An interruption template was computed by averaging the patterns for 10 TRs after the fifth TR from the interruption onset. Story-to-interruption inter-subject pattern correlation (ISPC) was the Pearson correlation between a participant's story template and the corresponding interruption template of the comparison group. We evaluated five inter-subject comparisons: three within-condition (IP-IP, SP-SP, IT-IT), where the comparison was the mean interruption template of the other participants in the same condition, and two across-condition (IP-IT, IT-IP), where it was the mean interruption template of the other condition's group. A positive correlation indicates that the story pattern is sustained into the interruption phase, whereas a zero or negative correlation indicates a transformed representational relationship.

Because story-to-interruption ISPC values are statistically dependent across participants through their shared group-average comparison pattern, significance was assessed using the same delete-one-subject jackknife and sign-flip permutation procedure as in the pattern reliability analysis. For participant *j*, the jackknife pseudo-value was computed as

$\tilde{\theta}_{j} = n \cdot\hat{\theta} - (n - 1) \cdot\hat{\theta}_{-j}$,

where $\hat{\theta}$ is the full-group story-to-interruption ISPC and $\hat{\theta}_{-j}$ is the same estimate recomputed after removing participant j. The resulting pseudo-values therefore provided an estimate for the participant-level contribution to the group ISPC. These participant-level estimates were submitted to a permutation test, with the null distribution built by independently sign-flipping the pseudo-value across participants over 10,000 iterations and recording the group mean. Because an inverted representational format predicts a negative correlation, significance was assessed using a one-sided lower-tail test, with P = (k + 1)/(N + 1), where k is the number of permuted means less than or equal to the observed group mean and N = 10,000.

***Story-to-interruption inversion selectivity analysis.*** Having observed a negative relationship between the story and interruption patterns, we next tested whether this inversion was specific to individual epochs. If the transformation is epoch-selective, story-to-interruption ISPC should be more negative for matching than for mismatching epoch pairs. Using the same story and interruption templates as described above, we computed story-to-interruption ISPC for every pair of epochs by correlating a participant's story template from each epoch with the comparison group's interruption template from every epoch. This yielded one matching correlation for the corresponding epoch pairs and one for mismatching epoch pairs. For participant j, inversion selectivity was defined as

Δ_j_ = r_match,j_ − r_mismatch,j_,

where r_match,j_ is the mean story-to-interruption ISPC for matching epoch pairs and r_mismatch,j_ is the mean across all mismatching epoch pairs. More negative values therefore indicate greater epoch-specific inversion. Group transformation selectivity was defined as the mean Δ_j_ across participants and was computed under the same five inter-subject comparisons as the transformation analysis (IP-IP, SP-SP, IT-IT, IP-IT, and IT-IP).

Because the ISPC values are statistically dependent, inversion selectivity was assessed using a within-participant epoch-label permutation test on the group-mean selectivity statistic (46), following the pattern selectivity analysis. Under the null hypothesis that the inversion contains no epoch-specific information, the matching and mismatching labels are exchangeable. For each of 10,000 permutations, each participant’s matching and mismatching ISPC values were pooled and randomly reassigned to sets of the original sizes and the matching minus mismatching selectivity statistic was recomputed, yielding a null distribution of group-mean selectivity values. Because epoch-specific inversion predicts a more negative correlation for matching than mismatching epochs, the observed selectivity was evaluated using a one-sided lower-tail permutation test, with P = (k + 1)/(N + 1), where k is the number of permuted means less than or equal to the observed group mean and N = 10,000. This procedure preserves the within-participant dependence structure of the ISPC values and examines whether the story-to-interruption inversion carries epoch-specific information.

***Brain-behavior modeling.*** We tested whether the shared PMC pattern sustained across the interruption and the hippocampal boundary response (defined in the Hippocampal boundary activity section above) predicted neural resumption following interruption and subsequent narrative memory across the 57 interrupted participants.

Neural resumption quantified how closely the neural patterns within predefined DMN ROIs (excluding PMC) realigned with those in the control group after participants returned to the story. At each return from interruption, an interrupted participant’s DMN pattern was correlated with the mean CT group pattern at the corresponding TRs, from 4 to 8 TRs after the return cue that marked the resumption of the story, to account for the immediate hemodynamic lag. The five correlations were averaged to obtain an epoch-level realignment score, and these scores were then averaged across epochs to obtain one neural realignment score per participant. Narrative memory was defined as the proportion of the 240 main-narrative sentences recalled by each participant during free spoken recall.

PMC pattern persistence quantified how stably the shared interruption pattern was sustained over the interruption phase. For each matching epoch, a participant's PMC pattern at each TR was correlated with the mean pattern of the other participants across all pairs of TRs in the 10-TR window beginning 5 TRs from interruption onset (TRs 5 to 14 after onset, 7.5 to 22.5 s), yielding a 10 x 10 time-time ISPC matrix. The 100 correlation values were averaged to obtain one persistence score per epoch, and these scores were then averaged across epochs to obtain one PMC pattern-persistence score per participant. Higher values therefore indicate greater temporal stability of the shared PMC pattern during interruption. Section S16 additionally checked increasingly distant off-diagonal values to quantify how stably the shared PMC pattern persisted over time.

We then used neural predictors to model our neural outcome variable (neural resumption) and our behavioral outcome variable (narrative memory). We fit ordinary-least-squares regression models pooling across all interrupted participants that included condition as a fixed effect to account for the mean differences between the IP, IT, SP groups. Three models were fit for each outcome: the shared PMC pattern persistence, the hippocampal boundary activity, and both predictors together. For each predictor we reported the unstandardized regression coefficient, its standard error, the t statistic, and the two-sided p value. The resumption models were fit on all 57 interrupted participants and the recall models on the 55 participants with a valid free-recall score.

***Statistical reporting***

Unless explicitly specified as one-sided in the analysis definitions above, statistical tests were two-sided. For parametric tests, we reported the test statistic and degrees of freedom; for permutation and sign-flip tests, we reported the number of participants. The resampling procedures described above determined statistical significance; 95% confidence intervals were reported to characterize the uncertainty of the corresponding effect estimates. The confidence interval is a parametric or bootstrap summary of the sampling distribution of the estimate, whereas each P value is the non-parametric test that determined its statistical significance reported. Confidence intervals for Fisher-z group means in the pattern-reliability and story-to-interruption transformation analyses were normal-approximation intervals (mean ± 1.96 SE across participants). Confidence intervals for matching-minus-mismatching selectivity estimates were percentile intervals derived from 10,000 participant bootstrap resamples. Confidence intervals for pattern-evolution estimates were t intervals on the participant-level slopes, using (n-1) degrees of freedom.

***Use of large language models***

Claude (Anthropic; Claude Opus 4.8, Claude Opus 5, Claude Fable 5, and Claude Fable 5.1, used through Claude Code) assisted the authors with three tasks: consolidating the authors’ existing analysis and figure-generation code into a documented repository, with every analysis re-run and its outputs verified against the authors’ original results; debugging the plotting code that renders the author-designed schematic illustrations; and editorial checks of author-written text for spelling, grammar, and journal-format compliance. The models were directed by short author-written instructions in interactive sessions; no generative prompt produced any part of the study, and no generative-image model was used. Hypotheses, study design, data collection, analyses, statistical inference, interpretation, and writing are the work of the authors, who reviewed every model-assisted edit.

**Supplementary Text**

***S1. Whole-brain inter-subject correlation across the four conditions***

We first examined whether the narrative drove reliably shared cortical responses in every condition. Whole-brain inter-subject correlation (ISC) maps during story listening were computed across the Schaefer 400-parcel, 17-network atlas (*38*) for each condition, excluding the first 8 TRs of story segments following an interruption to minimize hemodynamic carry-over from the preceding interruption. Parcel-level ISC was tested against zero across participants with Benjamini-Hochberg FDR correction across the 400 parcels.

The narrative elicited significant ISC (FDR < 0.05) across most of cortex in every condition: 345 of 400 parcels in continuous (CT), 379 in intact-pause (IP), 359 in intact-ToM (IT), and 336 in scrambled-pause (SP). Synchrony was strongest in auditory and lateral superior temporal cortex and extended into higher-order association areas, including default-mode, frontoparietal-control, salience/ventral-attention and temporoparietal networks, with broadly overlapping topographies across the four conditions (Fig. S3). Thus, despite the differences in narrative structure and interruption content across conditions, all groups showed widespread shared neural responses during story listening, providing a common basis for the subsequent multivoxel-pattern analysis.

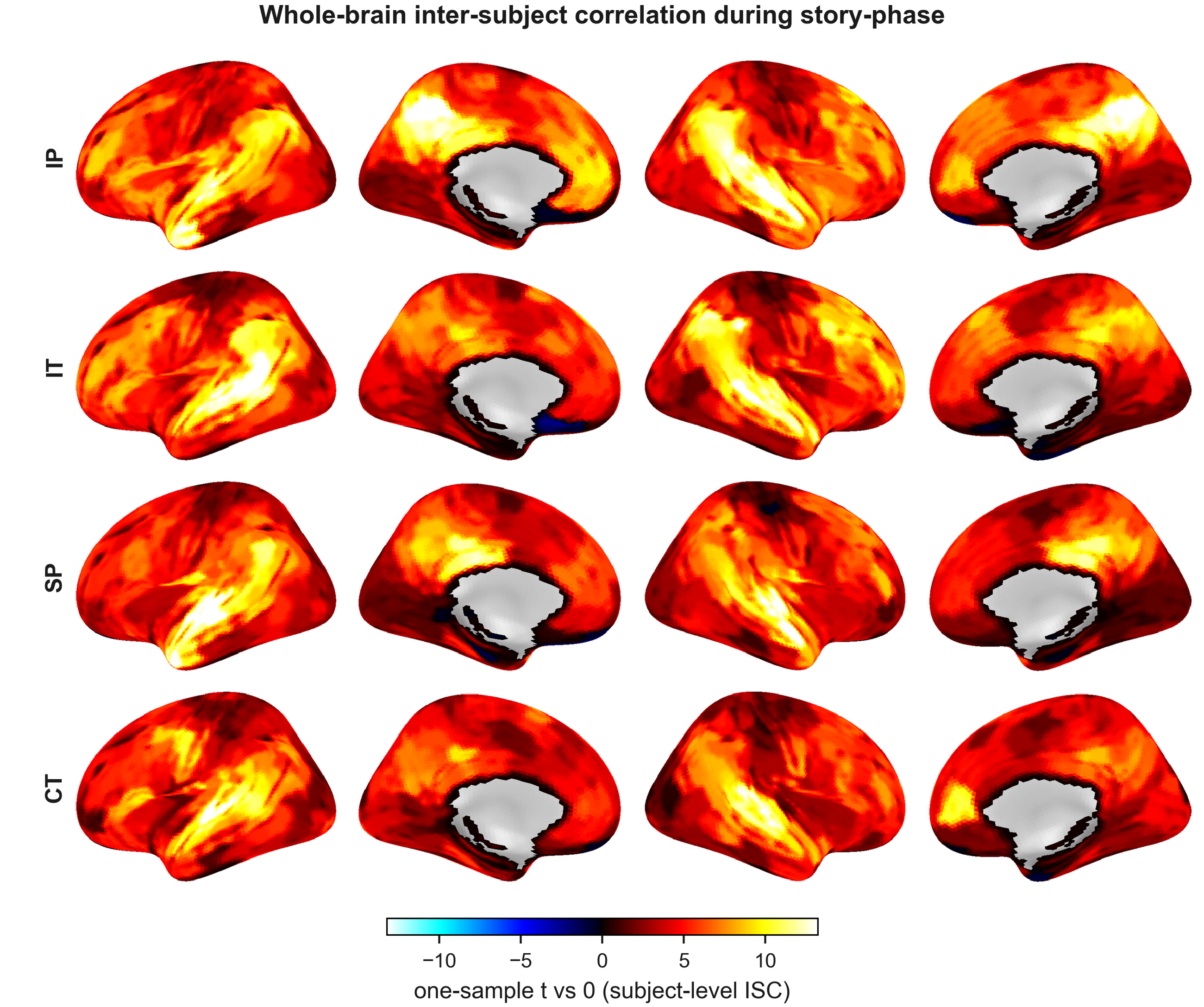

**Fig. S3.** Whole-brain inter-subject correlation (ISC) during story listening by condition. Per-parcel one-sample t statistic (participant-level ISC tested against zero) for the intact-pause (IP; n = 19), intact-ToM (IT; n = 19), scrambled-pause (SP; n = 19), and continuous (CT; n = 16) conditions. Significant ISC (Benjamini-Hochberg FDR < 0.05, across the 400 Schaefer parcels) was widespread in every condition, with the strongest synchrony in auditory and superior temporal cortex and extending into higher-order association cortex, including the default-mode network.

***S2. Whole-brain interruption-onset response***

We next examined the whole-brain response at interruption onset, to extend the hippocampal boundary response reported in the main text to the entire cortex. The same post-minus-pre boundary response was computed across the Schaefer 400-parcel 17-network atlas (*38*) for each interruption condition. For each parcel and participant, activity was averaged across the 5 TRs following the onset and contrasted with the 5 TRs immediately preceding onset, then averaged across epochs. Parcel-level boundary responses were tested against zero across participants using a two-sided one-sample t-test, with Benjamini-Hochberg FDR correction across the 400 parcels.

Interruption onset elicited widespread cortical responses in all three interrupted conditions, with significant responses (FDR < 0.05) in 277 of 400 parcels in intact-pause (IP), 204 in intact-ToM (IT), and 141 in scrambled-pause (SP). Activity decreased in auditory and language processing areas under the pause conditions and increased in salience/ventral-attention, frontoparietal-control cortex, and parts of the DMN across conditions (Fig. S4A). In the PMC, activity increased immediately after each interruption onset across the 57 interrupted participants [mean post-minus-pre response = 0.139, 95% CI (0.086, 0.192); t(56) = 5.26, P < 0.001]; under IT, PMC activity subsequently fell below baseline during the theory-of-mind task, whereas under IP and SP it returned toward baseline without a sustained decrease (Fig. S4B). Thus, interruption onset produced a widespread, network-organized cortical state change, providing the broader context for the sustained PMC pattern effects examined in the subsequent analyses.

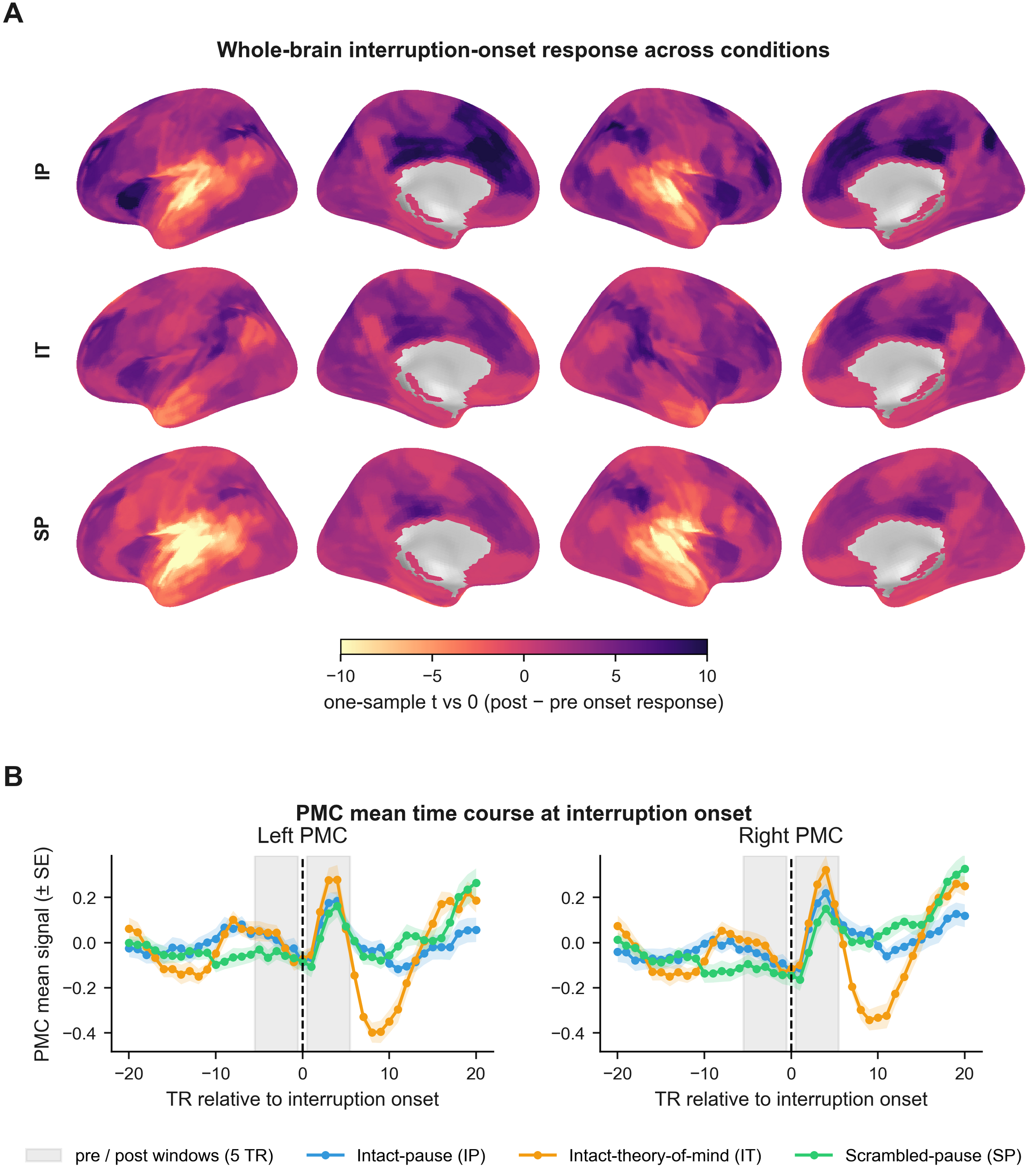

**Fig. S4.** Whole-brain interruption-onset response by condition. (A) Per-parcel one-sample t-map of the interruption onset response for the intact-pause (IP), intact-theory-of-mind (IT), and scrambled-pause (SP) conditions (n = 19 each). Significant responses (Benjamini-Hochberg FDR < 0.05, across the 400 Schaefer parcels) were widespread, including decreases in auditory and language areas and increases across higher-order association cortex. (B) Left and right PMC activity around interruption onset for each condition: signal was first z-scored based on within-subject story-phase activity, averaged across all epochs and subjects per condition, and shown as the group mean ± SEM across participants; gray bands indicate the 5-TR pre-onset and post-onset windows that were contrasted to compute the area’s interruption boundary activity.

***S3. Interruption-pattern reliability across the pre-selected regions of interest***

We first examined whether the interruption-phase multivoxel pattern was reliably shared across participants in each pre-selected ROI. Patterns were measured over the 10-TR interruption window beginning 5 TRs after onset. Pattern reliability was tested via three within-condition inter-subject comparisons (IP-IP, SP-SP, IT-IT) and one across-condition comparison (IT-IP), in which each intact-ToM participant was compared with the IP group mean to test whether the interruption pattern was preserved when attention was diverted by the theory-of-mind task.

Within each interruption condition, the interruption pattern was reliably shared across participants in every pre-selected ROI (all sign-flip P ≤ 0.006, n = 19 per condition; Fig. S5). Across conditions, however, IT participants showed a reliably positive match to the IP pattern in only three of the default-mode regions: PCC [IT-IP: Fisher-z = 0.104, 95% CI (0.079, 0.130), P < 0.001, n = 19], dmPFC [IT-IP: Fisher-z = 0.040, 95% CI (0.017, 0.063), P = 0.002, n = 19], and PMC [IT-IP: Fisher-z = 0.046, 95% CI (0.026, 0.067), P < 0.001, n = 19]. In contrast, the IT-IP pattern correlation was negative in the sensory and language regions [A1+: Fisher-z = −0.113, 95% CI (−0.134, −0.092); mSTG: Fisher-z = −0.127, 95% CI (−0.149, −0.106); n = 19 each], consistent with the different sensory input during the IT and IP interruptions. Thus, reliable interruption patterns were widespread within conditions, whereas preservation of the pattern across IP and IT was restricted to a subset of higher-order regions.

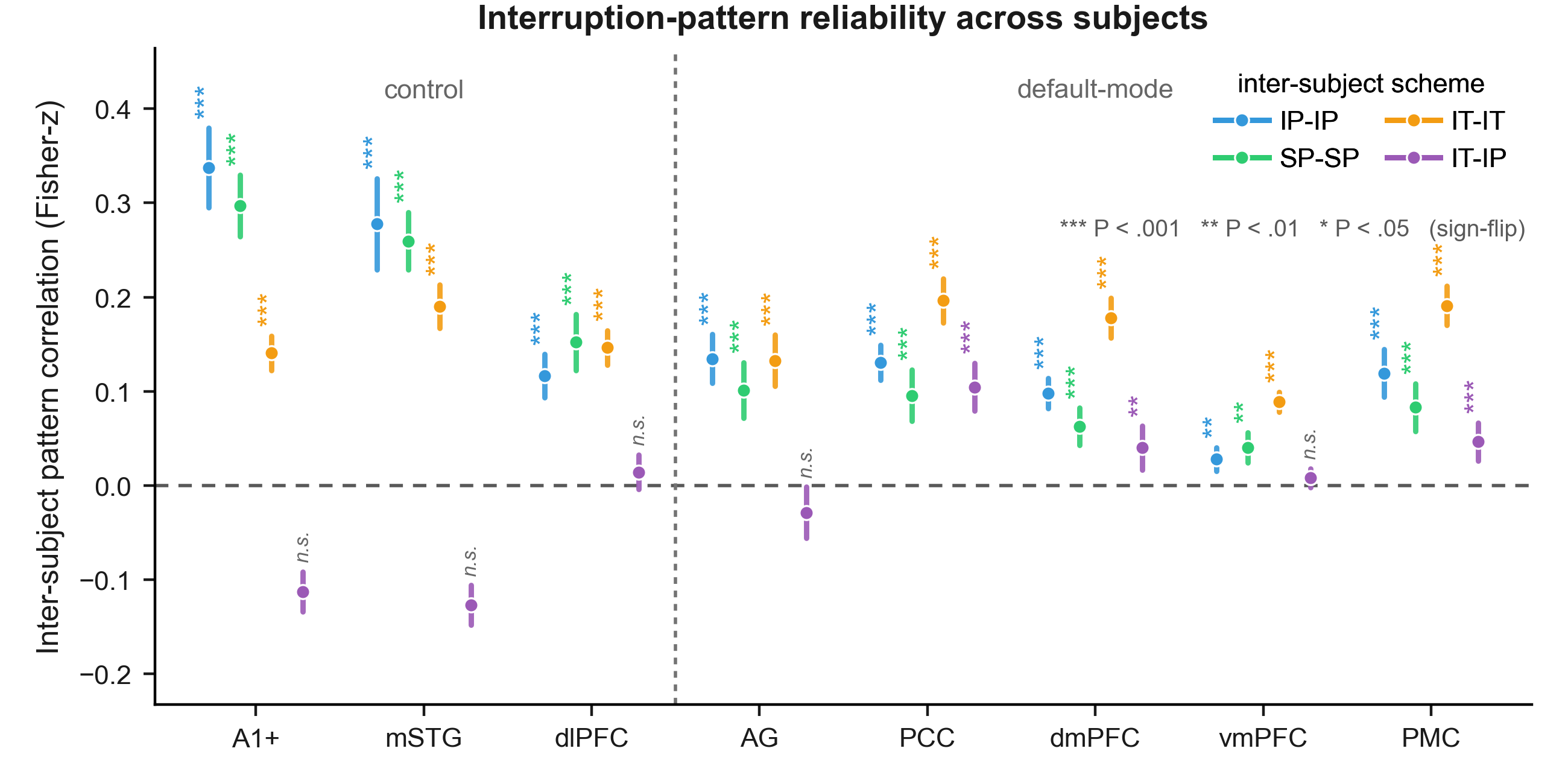

**Fig. S5.** Pattern reliability during the interruption phase across pre-selected regions of interest. Fisher-z group-mean ISPC with 95% CI for the within intact-pause (IP-IP), within scrambled-pause (SP-SP), within intact-ToM (IT-IT), and across-condition intact-ToM to intact-pause (IT-IP) comparisons (n = 19 per comparison). The vertical dotted line separates control ROIs from the predefined DMN ROIs; the horizontal dotted line marks zero. Asterisks indicate statistical significance of the one-sided sign-flip reliability test: ***=P < 0.001, **=P < 0.01, *=P < 0.05; n.s., not significant. Because the test is one-sided for positive reliability, negative correlations such as A1+ in the IT-IP comparison are marked n.s. even though these correlations are reliably negative.

***S4. Shared interruption pattern differed between the scrambled and intact narratives***

The interruption-phase PMC pattern was reliably shared and epoch-specific within both intact-pause (IP) and scrambled-pause (SP), although selectivity was weaker under SP. We next asked whether the same epoch-specific pattern was shared across IP and SP. Because the two groups heard the same story segments in different orders, SP interruptions could be aligned to IP interruptions in two ways. Alignment by serial position (SP-IP) paired interruptions occurring at the same serial position in the run but following different story segments. Re-ordering SP interruptions into the intact narrative sequence (unscrambled SP-IP) instead paired interruptions following the same story segment, thereby matching the immediately preceding story content.

These two alignments tested different explanations for the interruption pattern. If it reflected only a temporally evolving state associated with progression through the run, cross-condition epoch-selectivity should be significant when SP and IP were aligned by serial position. If it reflected only the immediately preceding story segment, cross-condition epoch-selectivity should be significant after aligning SP epochs to their original (un-scrambled) positions in the intact narrative. Results showed that selectivity was not significant when epochs were aligned by serial position [SP-IP: selectivity = 0.006, 95% CI (−0.006, 0.018), epoch-label permutation P = 0.247, n = 19] or after reordering SP epochs to match the IP epoch with the same bordering story content [SP-IP-unscrambled: selectivity = −0.001, 95% CI (−0.015, 0.015), P = 0.526, n = 19]. IP within-condition selectivity exceeded both across-condition estimates [Welch two-sided t(36.0) = 3.17, P = 0.003, d = 1.03; t(34.4) = 3.44, P = 0.002, d = 1.12]. Thus, the epoch-specific interruption pattern observed in IP depended on intact narrative organization rather than only on temporal position in the run or the local story segment preceding each interruption.

***S5. Pre-selected control and default-mode regions: only PMC exhibited all three signatures of an evolving background process***

To examine whether the interruption pattern signatures observed in PMC extended to other pre-selected DMN areas, we applied the same selectivity and pattern-evolution analyses to ROIs with reliable interruption ISPC. Analyses were conducted for three within-condition schemes (IP-IP, SP-SP, IT-IT), and one across-condition scheme (IT-IP). The IP-IP, SP-SP, and IT-IP comparisons were the primary tests of the proposed background representation, with IT-IP testing whether the interruption pattern was preserved when attention was diverted by the story-unrelated theory-of-mind task.

Among regions with reliable interruption ISPC (reported in S3), three of the four remaining default-mode regions carried epoch-specific patterns in the IP condition: PCC [IP-IP: selectivity = 0.016, 95% CI (0.009, 0.022), epoch-label permutation P = 0.030, n = 19], dmPFC [IP-IP: selectivity = 0.014, 95% CI (0.007, 0.021), P = 0.026, n = 19], and vmPFC [IP-IP: selectivity = 0.013, 95% CI (−3.4 × 10^−5^, 0.025), P = 0.026, n = 19]. In the SP condition, PCC [SP-SP: selectivity = 0.015, 95% CI (0.008, 0.023), P = 0.026, n = 19] and vmPFC [SP-SP: selectivity = 0.021, 95% CI (0.010, 0.033), P = 7 × 10^−4^, n = 19] stayed epoch specific. Under the competing theory-of-mind task in the IT condition, PCC was the only additional DMN area whose pattern matched the IP group in an epoch-selective manner [IT-IP: selectivity = 0.020, 95% CI (0.014, 0.025), P = 0.010, n = 19].

Pattern evolution was then examined in regions with reliable and epoch-selective interruption patterns. None of the remaining DMN areas showed a significant decline in pattern similarity with epoch-distance under IP [IP-IP: PCC, dmPFC and vmPFC, all P > 0.1, n = 19 each]. Under SP, no region showed a systematic decline with epoch distance, consistent with the absence of coherent context to integrate across epochs. Under IT-IP, PCC likewise showed no significant decline (IT-IP: P = 0.101, n = 19).

Therefore, PMC was the only pre-selected DMN region to show all three interruption-pattern neural signatures predicted of the background process: reliability, epoch-selectivity, and systematic evolution across the intact narrative. PCC showed reliable and epoch-selective interruption patterns but no gradual change with epoch distance. AG showed the complementary pattern: when tested outside the sequential procedure its pattern similarity declined with epoch distance, but it did not exhibit an epoch-selective interruption pattern. Together, these dissociations indicate that the full combination of interruption-pattern signatures was specific to PMC among the pre-selected DMN areas.

**Table S2.** Summary table for interruption-pattern signatures across the pre-selected default-mode and control regions. Summary of pattern reliability, epoch-selectivity, and pattern evolution across the five pre-selected default-mode regions (AG, PCC, dmPFC, vmPFC, and PMC) and three control regions (A1+, mSTG, dlPFC). The first nine columns show the three signatures tested via IP-IP, SP-SP, and IT-IP comparisons. The final three columns show IT-IT, which reflects patterns evoked by the theory-of-mind interruption stimuli and is included for reference. Green check marks indicate that the predicted criterion was met (P < 0.05); red crosses indicate that the criterion was tested but not met; gray cells indicate analyses were not conducted under the sequential testing procedure because a preceding criterion was not met. Note that we predicted an absence of systematic narrative-scale pattern evolution for the SP-SP comparisons.

| ROI | Reliable | | | Selective | | | Evolve | | | IT-IT | | |
| --- | --- | --- | --- | --- | --- | --- | --- | --- | --- | --- | --- | --- |
|  | IP-IP | SP-SP | IT-IP | IP-IP | SP-SP | IT-IP | IP-IP | SP-SP | IT-IP | Reliable | Selective | Evolve |
| AG | **✓** | **✓** | **✗** | **✗** | **✗** | **✗** | **✓** | **✓** | **✓** | **✓** | **✓** | **✗** |
| PCC | **✓** | **✓** | **✓** | **✓** | **✓** | **✓** | **✗** | **✓** | **✗** | **✓** | **✓** | **✗** |
| dmPFC | **✓** | **✓** | **✓** | **✓** | **✗** | **✗** | **✗** | **✓** | **✗** | **✓** | **✓** | **✗** |
| vmPFC | **✓** | **✓** | **✗** | **✓** | **✓** | **✓** | **✗** | **✓** | **✗** | **✓** | **✓** | **✗** |
| PMC | **✓** | **✓** | **✓** | **✓** | **✓** | **✓** | **✓** | **✓** | **✓** | **✓** | **✓** | **✓** |
| A1+ | **✓** | **✓** | **✗** | **✗** | **✗** | **✗** | **✓** | **✓** | **✓** | **✓** | **✓** | **✗** |
| mSTG | **✓** | **✓** | **✗** | **✗** | **✗** | **✗** | **✗** | **✓** | **✓** | **✓** | **✓** | **✓** |
| dlPFC | **✓** | **✓** | **✗** | **✗** | **✗** | **✗** | **✗** | **✓** | **✓** | **✓** | **✓** | **✓** |

We applied the same analyses to the three pre-selected control regions targeting auditory, language and working-memory processing areas: early auditory cortex (A1+), middle superior temporal gyrus (mSTG), and dorsolateral prefrontal cortex (dlPFC). These areas were not expected to carry epoch-selective interruption patterns in the IP-IP, SP-SP, IT-IP tests, but were expected to show epoch-selective interruption patterns in the IT-IT, driven by the distinct theory-of-mind stimuli presented during each interruption. Consistent with these predictions, interruption patterns were reliable within IP-IP and SP-SP (all *P* < 1 × 10^−4^), but none of these control regions were epoch-specific in either condition (all *P* > 0.1). In the IT condition, A1+ and mSTG story patterns were negatively correlated with the IP interruption pattern [Fisher-z = −0.113, 95% CI (−0.134, −0.092) and −0.127, 95% CI (−0.149, −0.106)], while the dlPFC story patterns were not correlated with the IP interruption patterns (P = 0.072). Therefore, no ROIs in the control areas advanced to the selectivity or pattern-evolution tests for the IT-IP comparison. In contrast, all three ROIs showed reliable and epoch-selective patterns for IT-IT comparisons, as expected for regions generating distinct responses to the various theory-of-mind vignettes.

**Table S3.** Interruption-pattern statistics across the pre-selected cortical ROIs. For each ROI in control and default-mode networks, the table reports interruption-pattern reliability, epoch-selectivity, and pattern evolution via the IP-IP, SP-SP, and IT-IP comparisons (when the assumption is met to perform the test). Cells marked N/A were not tested because a preceding criterion in the sequential testing procedure was not met (Table S2). Asterisks indicate permutation P values: *=P < 0.05, **=P < 0.01, ***=P < 0.001.

| Cond | ROI | Mean ISPC (Z) | 95% CI | Selectivity Δ | Perm P | Evolve slope | Slope P |
| --- | --- | --- | --- | --- | --- | --- | --- |
| IP-IP | AG | 0.134 | (0.109, 0.160) | 0.005 | .313 | N/A | N/A |
|  | PCC | 0.131 | (0.112, 0.149) | 0.016 | .030^*^ | −0.0009 | .188 |
|  | dmPFC | 0.098 | (0.082, 0.114) | 0.014 | .026^*^ | −0.0004 | .535 |
|  | vmPFC | 0.028 | (0.015, 0.040) | 0.013 | .026^*^ | 0.0009 | .368 |
|  | PMC | 0.119 | (0.094, 0.145) | 0.034 | <0.001^***^ | −0.0036 | <0.001^***^ |
|  | A1+ | 0.337 | (0.295, 0.380) | 0.002 | .357 | N/A | N/A |
|  | mSTG | 0.278 | (0.229, 0.326) | 0.005 | .251 | N/A | N/A |
|  | dlPFC | 0.116 | (0.093, 0.139) | 0.004 | .248 | N/A | N/A |
| SP-SP | AG | 0.101 | (0.072, 0.131) | 0.008 | .195 | N/A | N/A |
|  | PCC | 0.096 | (0.068, 0.123) | 0.015 | .026^*^ | 0.0005 | .553 |
|  | dmPFC | 0.062 | (0.043, 0.082) | 0.005 | .197 | N/A | N/A |
|  | vmPFC | 0.040 | (0.024, 0.056) | 0.021 | <0.001^***^ | 0.0002 | .785 |
|  | PMC | 0.083 | (0.058, 0.108) | 0.015 | .027^*^ | 0.0000 | .989 |
|  | A1+ | 0.297 | (0.265, 0.330) | 0.003 | .320 | N/A | N/A |
|  | mSTG | 0.260 | (0.229, 0.290) | 0.005 | .266 | N/A | N/A |
|  | dlPFC | 0.152 | (0.122, 0.182) | 0.008 | .138 | N/A | N/A |
| IT-IP | AG | −0.029 | (−0.056, −0.002) | N/A | N/A | N/A | N/A |
|  | PCC | 0.104 | (0.079, 0.130) | 0.020 | .0098** | −0.0011 | .101 |
|  | dmPFC | 0.040 | (0.017, 0.063) | 0.010 | .065 | N/A | N/A |
|  | vmPFC | 0.008 | (−0.002, 0.018) | N/A | N/A | N/A | N/A |
|  | PMC | 0.046 | (0.026, 0.067) | 0.021 | .003^**^ | −0.0022 | .005^**^ |
|  | A1+ | −0.113 | (−0.134, −0.092) | N/A | N/A | N/A | N/A |
|  | mSTG | −0.127 | (−0.149, −0.106) | N/A | N/A | N/A | N/A |
|  | dlPFC | 0.014 | (−0.004, 0.032) | N/A | N/A | N/A | N/A |

**Table S4.** Complete statistical parameters for the analyses reported in the main text. For each main-text statistical claim: effect estimate, 95% confidence interval (CI), standard error (SE), test statistic with degrees of freedom (or effect size where the primary test is non-parametric), P value, and sample size; the main text reports the abbreviated form (estimate, test statistic, P). Selectivity and pattern-evolution P values are within-participant permutation tests; story-to-interruption P values are jackknife sign-flip tests; regression rows are ordinary-least-squares models pooled across the interrupted conditions with condition entered as a fixed effect. Confidence intervals were computed as defined in the Statistical reporting section (normal-approximation intervals for Fisher-z group means, participant-bootstrap percentile intervals for selectivity estimates, and t intervals for pattern-evolution slopes); confidence intervals for the regression coefficients are t-based intervals (b ± t0.975 × SE); dz, Cohen's dz; d, Cohen's d; diff., group difference. Values reproduce the archived analysis outputs.

| Analysis | Comparison / model | Estimate | 95% CI | SE | Test statistic | P | n |
| --- | --- | --- | --- | --- | --- | --- | --- |
| Recall (behavior) | Omnibus, one-way ANOVA | — | — | — | F(3, 67) = 5.95 | 0.001 | 71 |
|  | SP vs CT (Welch, Bonferroni) | — | — | — | t(25.4) = 3.35 | 0.015 | 34 |
|  | SP vs IP (Welch, Bonferroni) | — | — | — | t(29.9) = 4.39 | < 0.001 | 37 |
| Comprehension (behavior) | Omnibus, one-way ANOVA | — | — | — | F(3, 55) = 8.01 | < 0.001 | 59 |
|  | SP vs CT (Welch, Bonferroni) | — | — | — | t(16.5) = 5.09 | < 0.001 | 23 |
|  | SP vs IP (Welch, Bonferroni) | — | — | — | t(32.0) = 4.05 | 0.002 | 34 |
|  | SP vs IT (Welch, Bonferroni) | — | — | — | t(31.9) = 3.35 | 0.013 | 34 |
| Hippocampal boundary response | post − pre, pooled IP+IT+SP; one-sample t | 0.049 | (0.018, 0.080) | 0.015 | t(56) = 3.20 | 0.002 | 57 |
| PMC epoch selectivity | IP-IP; epoch-label permutation | 0.034 | (0.022, 0.046) | 0.006 | dz = 1.24 | 2 × 10−4 | 19 |
|  | SP-SP | 0.015 | (0.005, 0.025) | 0.005 | dz = 0.68 | 0.027 | 19 |
|  | IT-IP | 0.021 | (0.013, 0.030) | 0.004 | dz = 1.15 | 0.003 | 19 |
|  | IP-IP vs SP-SP (Welch) | 0.019 (diff.) | — | — | t(34.4) = 2.36, d = 0.76 | 0.024 | 38 |
| PMC pattern-evolution slope | IP-IP; distance-label permutation (two-sided) | −3.6 × 10−3 | (−6.1 × 10−3, −1.1 × 10−3) | 1.2 × 10−3 | — | 9 × 10−4 | 19 |
|  | SP-SP | 1.3 × 10−5 | (−2.8 × 10−3, 2.8 × 10−3) | 1.3 × 10−3 | — | 0.989 | 19 |
|  | SP-SP-unscrambled | 6.6 × 10−5 | (−2.9 × 10−3, 3.1 × 10−3) | 1.4 × 10−3 | — | 0.943 | 19 |
|  | IT-IP | −2.2 × 10−3 | (−4.4 × 10−3, −2.0 × 10−5) | 1.0 × 10−3 | — | 0.005 | 19 |
| Story-to-interruption correlation (Fisher-z) | IP-IP; jackknife sign-flip (one-sided) | −0.245 | (−0.291, −0.199) | — | — | < 1 × 10−4 | 19 |
|  | SP-SP | −0.209 | (−0.262, −0.156) | — | — | < 1 × 10−4 | 19 |
|  | IT-IT | −0.171 | (−0.210, −0.133) | — | — | 2 × 10−4 | 19 |
| Story-to-interruption selectivity (Δ) | IP-IP; epoch-label permutation (one-sided) | −0.064 | (−0.083, −0.042) | — | dz = −1.36 | 0.001 | 19 |
|  | SP-SP | −0.038 | (−0.057, −0.020) | — | dz = −0.90 | 0.024 | 19 |
|  | IT-IT | −0.058 | (−0.082, −0.036) | — | dz = −1.12 | 0.001 | 19 |
| Neural resumption (OLS, condition fixed effect) | PMC persistence alone | 0.386 | (0.231, 0.541) | 0.077 | t(53) = 4.99 | < 0.001 | 57 |
|  | Hippocampal boundary alone | 0.093 | (0.029, 0.157) | 0.032 | t(53) = 2.91 | 0.005 | 57 |
|  | PMC persistence, joint model | 0.350 | (0.200, 0.501) | 0.075 | t(52) = 4.67 | < 0.001 | 57 |
|  | Hippocampal boundary, joint model | 0.069 | (0.014, 0.124) | 0.028 | t(52) = 2.50 | 0.016 | 57 |
| Narrative recall (OLS, condition fixed effect) | PMC persistence alone | 0.764 | (0.252, 1.275) | 0.255 | t(51) = 3.00 | 0.004 | 55 |
|  | Hippocampal boundary alone | 0.232 | (0.043, 0.421) | 0.094 | t(51) = 2.47 | 0.017 | 55 |
|  | PMC persistence, joint model | 0.663 | (0.156, 1.169) | 0.252 | t(50) = 2.63 | 0.011 | 55 |
|  | Hippocampal boundary, joint model | 0.185 | (0.003, 0.367) | 0.091 | t(50) = 2.04 | 0.047 | 55 |

***S6. Whole-brain Schaefer-400 reliability, selectivity, and pattern evolution***

Having observed three signatures predicted of a background process in PMC (among the pre-selected ROIs), we examined how these effects were distributed across cortex. We repeated the reliability, selectivity, and pattern-evolution analyses in each parcel of the Schaefer 400-parcel atlas for the four inter-subject comparisons (IP-IP, SP-SP, IT-IT, and IT-IP). Statistical inference followed the procedure applied to the pre-selected ROI, with parcel-level P values corrected across the 400 parcels by Benjamini-Hochberg FDR.

The interruption-phase pattern was reliable in 246 of 400 parcels in the IP-IP condition, 226 in the SP-SP condition, 379 in the IT-IT condition (where interruptions were filled with vignettes), and 79 when comparing across the IT and IP conditions. However, the interruption patterns were only epoch specific (i.e. statistically significantly selective) for 11 parcels in the IP-IP condition, 3 in the SP condition, and none when contrasting IT-IP and applying FDR correction. The 11 IP-IP parcels were scattered across default-mode, frontoparietal-control, salience/ventral-attention, somatomotor, visual and dorsal-attention networks rather than concentrated in any one network. The three SP-SP parcels lay in left somatomotor cortex, the right temporal pole, and right ventrolateral prefrontal cortex. As expected, IT-IT selectivity was widespread (348 parcels distributed across all cortical systems), consistent with the distinct theory-of-mind stimulus presented during each interruption. Under IP-IP, pattern evolution survived FDR correction in 5 parcels (bilateral precuneus, extrastriate visual cortex, and right inferior parietal cortex), whereas no parcel showed a significant decline under SP-SP.

In order to provide a view of the spatial distribution of effects, while not limiting only to FDR-corrected significant regions, Fig. S6 shows the spatial distributions of selectivity and pattern evolution at an uncorrected P < 0.1 threshold.

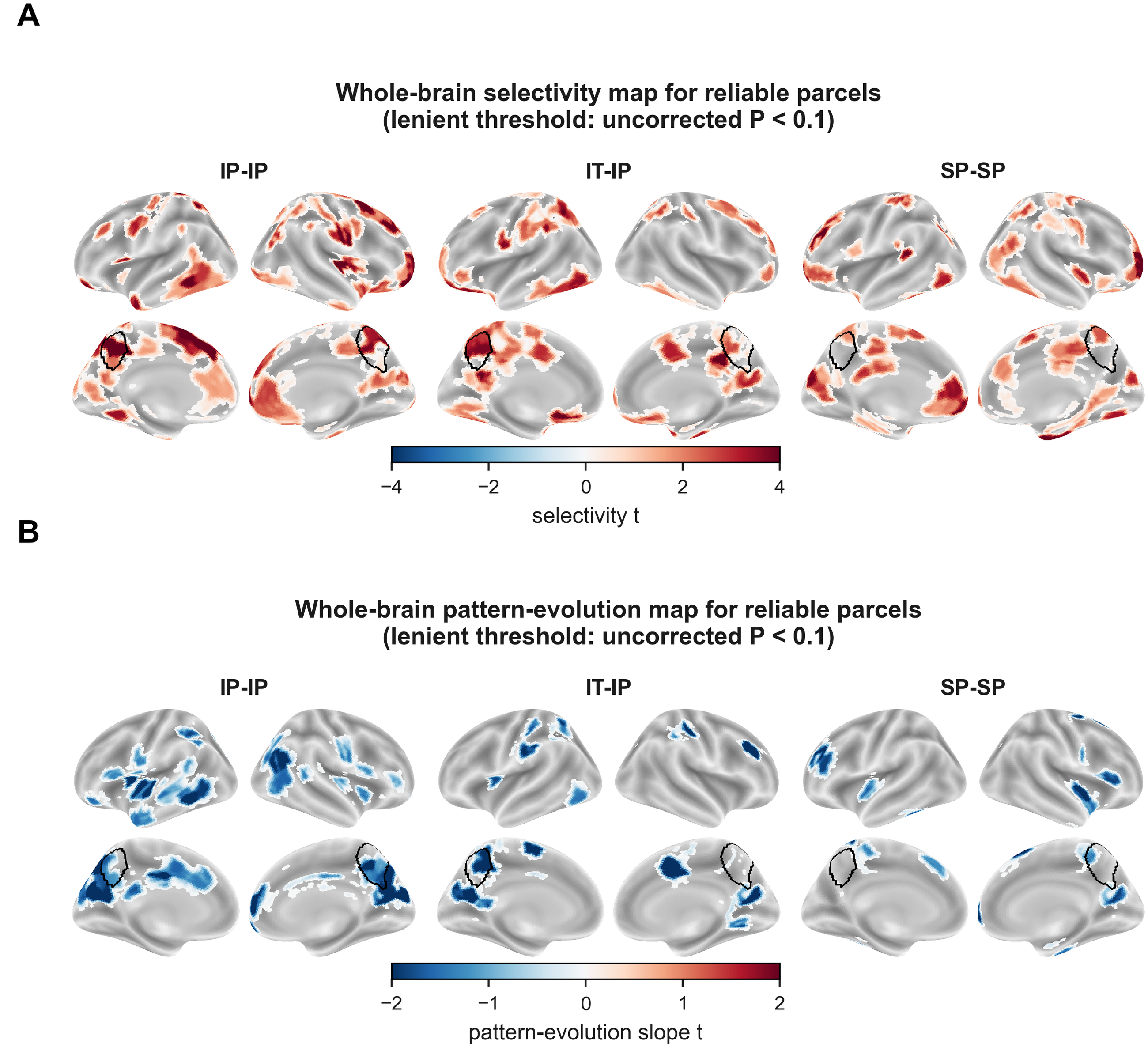

**Fig. S6.** Whole-brain Schaefer-400 selectivity and pattern evolution maps. (A) Per-parcel t-map of pattern selectivity (matching > mismatching) for the IP-IP, IT-IP, and SP-SP comparisons (n = 19 each). (B) Per-parcel t-map of pattern evolution (negative epoch-distance slope) for the IP-IP, IT-IP, and SP-SP comparisons. All maps show every parcel that was reliable at an uncorrected sign-flip P < 0.05 and reached an uncorrected permutation P < 0.10 in the hypothesized direction; the posterior medial cortex (PMC) is outlined in black. Both panels deliberately use this lenient display threshold to show the spatial distribution of effects; under the null, roughly 10% of the tested reliable parcels in each map (approximately 8 to 38 parcels per comparison) would be expected to pass by chance.

***S7. The PMC interruption-phase signatures generalized to a second narrative***

We next asked whether the PMC interruption-phase signatures generalized beyond the main narrative. Thus, we repeated the pattern reliability, selectivity, and pattern-evolution tests on a second narrative (the live-storytelling narrative, which comprised 12 segments and 11 interruption epochs). We employed the same dependent variable metrics, interruption window definitions, and statistical inference procedures as for the main narrative.

The PMC interruption pattern was reliably shared across participants in all four comparisons [Fisher-z group mean = 0.069 (IP-IP), 95% CI (0.050, 0.089); 0.058 (SP-SP), 95% CI (0.045, 0.071); 0.165 (IT-IT), 95% CI (0.136, 0.193); and 0.021 (IT-IP), 95% CI (0.004, 0.038); all sign-flip P ≤ 0.015, n = 19 each].

The PMC pattern was epoch-selective in three of the four comparisons [IP-IP: Δ = 0.021, 95% CI (0.005, 0.036), P = 0.009; IT-IT: Δ = 0.076, 95% CI (0.063, 0.090), P < 0.001; IT-IP: Δ = 0.013, 95% CI (0.002, 0.025), P = 0.037; n = 19 each] but reached only trend level in the SP-SP comparison [Δ = 0.011, 95% CI (0.002, 0.021), P = 0.082, n = 19].

The shared PMC pattern similarity across epochs again decreased with epoch distance under IP-IP [b = −5.1 × 10^−3^, 95% CI (−1.03 × 10^−2^, 1.2 × 10^−4^), P = 0.033, n = 19] and IT-IP [b = −6.1 × 10^−3^, 95% CI (−1.02 × 10^−2^, −2.0 × 10^−3^), P = 0.006, n = 19], but not under SP-SP [b = −8.4 × 10^−5^, 95% CI (−4.5 × 10^−3^, 4.3 × 10^−3^), P = 0.968, n = 19], reproducing the result observed with the main narrative. As for the main narrative, re-ordering the scrambled-pause epochs into the intact narrative sequence did not restore a decline either [unscrambled SP-SP: b = 1.8 × 10^−3^, 95% CI (−2.0 × 10^−3^, 5.7 × 10^−3^), P = 0.332, n = 19] (Fig. S7).

Thus, the second narrative reproduced the diagnostic PMC profile observed in the main narrative: interruption patterns were reliable and epoch-selective under the intact narrative, epoch specificity was preserved across IP and IT, and pattern similarity declined with narrative distance under IP-IP and IT-IP but not under scrambling. SP-SP selectivity was positive but did not reach significance in the second narrative. The story-to-interruption transformation also generalized to this narrative (Section S8).

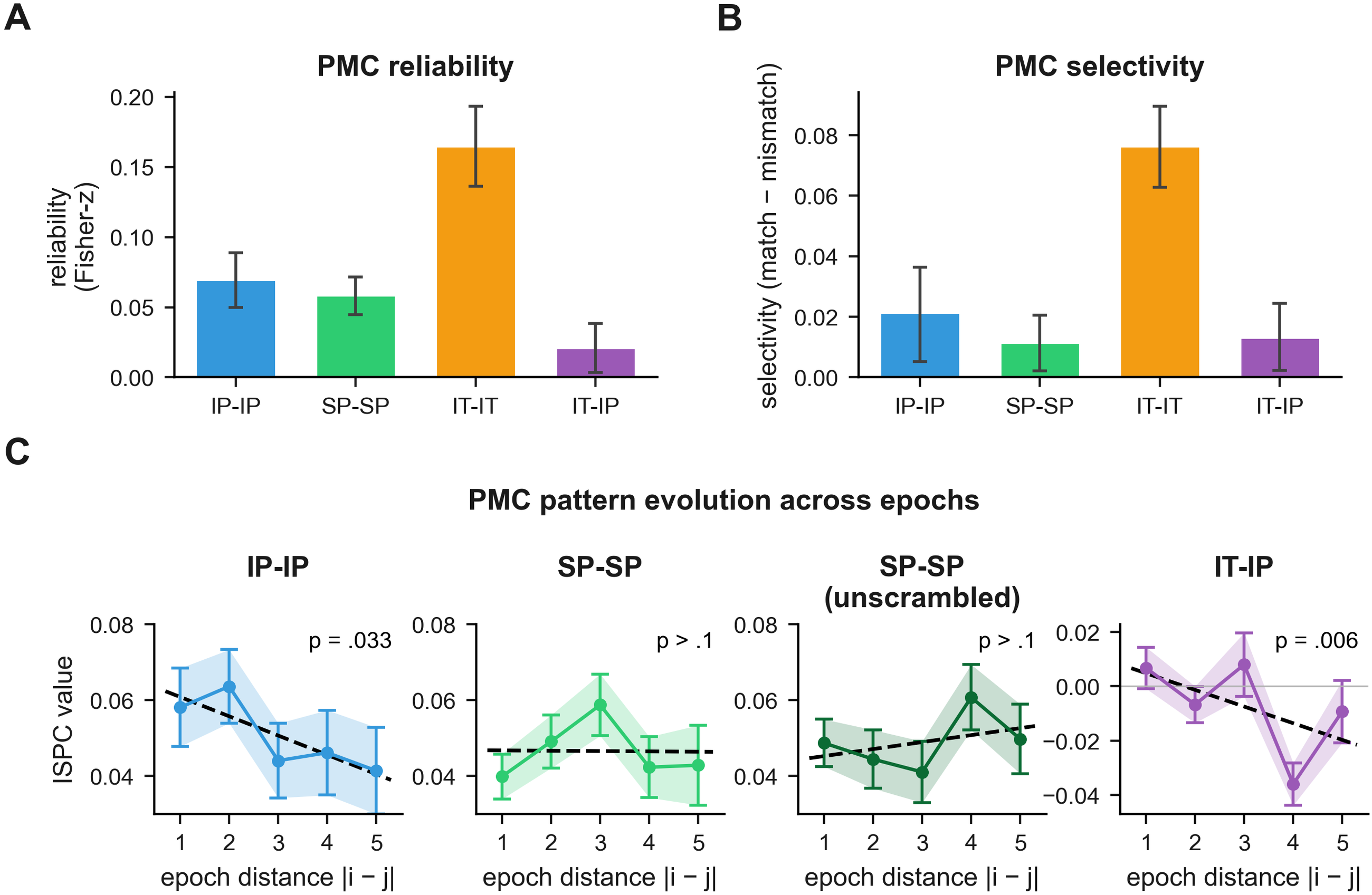

**Fig. S7. Generalization of the PMC interruption-phase pattern analyses to the live-storytelling narrative.** (A) Per-condition reliability (Fisher-z group-mean inter-subject pattern correlation) with 95% confidence intervals (n = 19 per comparison). (B) Per-condition selectivity (matching minus mismatching) with 95% confidence intervals. (C) Pattern similarity against epoch distance for the IP-IP, SP-SP, SP-SP (unscrambled), and IT-IP comparisons (shading shows ± SEM; the dashed line is the best-fitting slope). The live-storytelling narrative has 11 interruption epochs. The reliability, selectivity, and pattern-evolution effects generalize from the main narrative to this narrative.

***S8. The story-to-interruption transformation generalizes to a second narrative***

We next asked whether the story-to-interruption transformation effects generalized to the second narrative. The test for story-to-interruption transformation, the test for whether the transformation was epoch-selective, and the voxel-wise undershoot control analyses were repeated on the live-storytelling narrative (11 interruption epochs).

The transformation effect generalized to the second narrative. Each participant's PMC story-phase pattern was reliably negatively correlated with the comparison group's interruption-phase pattern in every comparison. The Fisher-z group means were −0.120 (IP-IP), 95% CI (−0.157, −0.082); −0.134 (SP-SP), 95% CI (−0.176, −0.091); −0.154 (IT-IT), 95% CI (−0.189, −0.119); −0.108 (IP-IT), 95% CI (−0.147, −0.070); and −0.088 (IT-IP), 95% CI (−0.124, −0.052); all sign-flip P ≤ 0.002, n = 19 each.

The transformation was also epoch-specific: the matching story-to-interruption correlation was more negative than the mismatching correlations in the SP-SP comparison [Δ = −0.045, 95% CI (−0.075, −0.014), P = 0.011], the IT-IT comparison [Δ = −0.092, 95% CI (−0.122, −0.058), P < 0.001], IP-IT [Δ = −0.079, 95% CI (−0.106, −0.052), P < 0.001], and the IT-IP comparison [Δ = −0.033, 95% CI (−0.055, −0.011), P = 0.035; n = 19 each]. The IP-IP effect was negative in the predicted direction but did not reach significance [Δ = −0.027, 95% CI (−0.054, 0.001), P = 0.071].

The undershoot control analyses again argued against a post-stimulus hemodynamic undershoot in PMC [PMC Q4 fraction = 0.04, 95% CI (0.001, 0.19), opposite to the undershoot direction], and in favor of something like undershoot in the auditory cortex [A1+ Q4 fraction = 1.00, bootstrap P < 0.001, n = 57 (19 per condition)] (Fig. S8). Thus, the transformation effect generalized across all five comparisons, while the epoch specificity was reproduced in four of the five comparisons. IP-IP comparisons showed the predicted negative correlation, but the effect did not reach statistical significance.

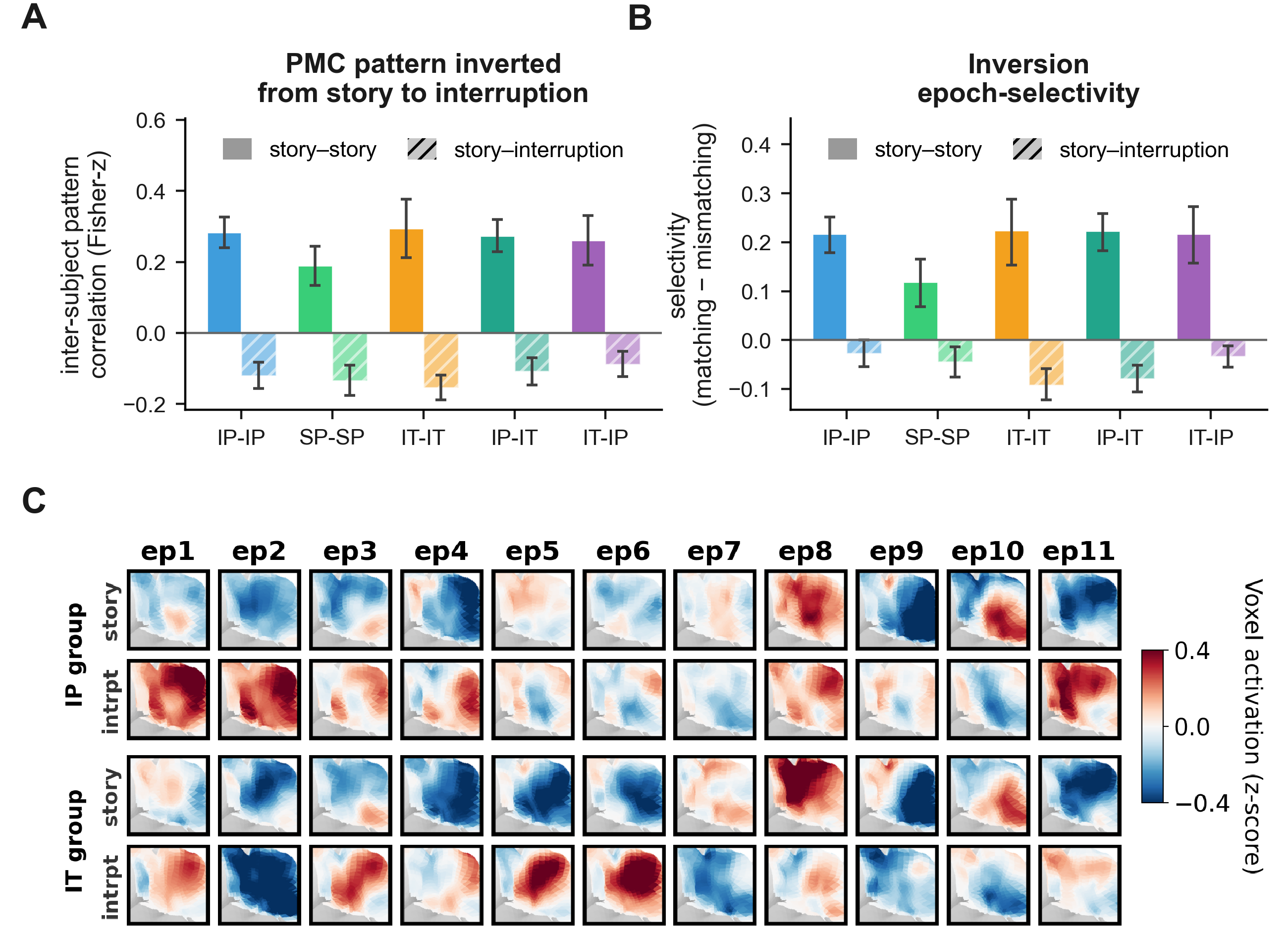

**Fig. S8. Generalization of the PMC transformation effects to a distinct narrative taken from a live storytelling event.** (A) Story-story and story-to-interruption inter-subject pattern correlations (Fisher-z group mean) across the five comparisons (n = 19 each); the negative story-to-interruption correlation (inversion) generalizes to the second narrative. (B) Inversion epoch-selectivity (matching more inverted than mismatching) across the comparisons. (C) The PMC multivoxel spatial patterns for the live-storytelling narrative: per-epoch story and interruption group-mean patterns for the IP and IT groups.

***S9. Story-to-interruption transformation: magnitude of the inversion***

The story-to-interruption correlation is measured in the presence of noise, so the magnitude of the correlation understates the proportion of reliable signal that is correlated or anticorrelated across phases. To quantify the extent of the inversion in PMC, we expressed the observed negative story-to-interruption correlation as a fraction of the largest negative correlation the data could support given the reliability of each phase: the negative geometric mean of the story-story and interruption-interruption reliabilities, where 0 corresponds to an orthogonal (unrelated) pattern and the ceiling to a complete sign-flip. We then corrected the correlation by that ceiling to estimate the noise-free correlation between the underlying story and interruption patterns, with a delete-one-participant jackknife 95% confidence interval. Specifically, the ceiling-corrected correlation was computed as $\rho_{true}=\frac{r_{SI}}{\sqrt{r_{SS}\cdot r_{II}}}$, where $r_{SI}$ is the observed story-to-interruption correlation and $r_{SS}$ and $r_{II}$ are the story-story and interruption-interruption reliabilities. The fraction of the distance to a complete inversion is $\frac{r_{SI}}{-\sqrt{r_{SS}\cdot r_{II}}}$, and the corresponding rotation angle is $\arccos(\rho_{true})$.

The observed PMC story-to-interruption correlation was negative and of a magnitude comparable to the positive story-story correlation on matched windows [IP-IP: story-to-interruption Fisher-z = −0.245, 95% CI (−0.291, −0.199); story-story Fisher-z = 0.347, 95% CI (0.288, 0.406); both sign-flip P < 0.001, n = 19]. This observed inversion captured a large fraction of the signal reliability ceiling [80.9% (IP-IP), 95.3% (SP-SP), and 59.9% (IT-IT)], and the ceiling-corrected correlation between the underlying patterns was strongly and reliably negative [ρ = −0.81, 95% CI (−0.94, −0.69) (IP-IP); −0.95, 95% CI (−1.00, −0.79), truncated at the −1 boundary (SP-SP); −0.60, 95% CI (−0.70, −0.46) (IT-IT)].

Interpreting these pattern changes as angles of vector rotation, the pattern angles [144° (IP-IP), 162° (SP-SP), 127° (IT-IT)] were all greater than the 90° orthogonal line and toward 180° (corresponding to an exact inversion or flip of the pattern vector). Thus, the inversion traveled most of the way to the ceiling set by the reliability of the data. This indicates a strong inversion of the story-phase pattern, rather than merely an orthogonalization.

The negative correlations between story and interruption phase decreased when we analyzed the data using a secondary preprocessing pipeline that lacked a high-pass temporal filter and nuisance regression. This could reflect (a) decreased signal quality with minimal preprocessing as well as (b) some of the inversion effect being magnified directly by the high-pass filter. We describe the transformation as an inversion because the pattern correlation (even in temporally unfiltered BOLD data) remained negative, and the pattern is clearly reliable, selective, and distinct from the story phase. However, more precisely characterizing the transformation will likely require triangulating with additional methods, such as intracranial recordings.

***S10. Story-to-interruption transformation: testing for hemodynamic-undershoot***

We can quantify the changes in BOLD signal from the story phase to the interruption phase by plotting the mean signals in each phase against one another. We can then divide the plot into quadrants (Q1: story positive to interruption positive; Q2: story negative to interruption positive; Q3: story negative to interruption negative; Q4 story positive to interruption negative).

A post-stimulus hemodynamic undershoot would drive voxels that were positive during the story below baseline during the interruption (a story-positive to interruption-negative flip, quadrant Q4) without the converse flip of story-negative voxels (quadrant Q2), so its signature is an excess of Q4 over Q2 inverting voxels. Thus, a pure post-stimulus undershoot predicts a Q4 fraction greater than the symmetric value of 0.5. We averaged each voxel's story-window and interruption-window value across all interruption epochs, pooled the three interruption conditions. We tested whether the resulting Q4 fraction exceeded 0.5 by computing a one-sided P value via a participant-wise bootstrap (5,000 resamples).

Primary auditory cortex and middle superior temporal gyrus showed the predicted undershoot asymmetry [Q4 fraction = 1.00, 95% CI (1.00, 1.00), bootstrap P < 0.001 for both, n = 57 (19 per condition)]. PMC did not show this asymmetry: its inverting voxels were approximately balanced between the two directions [Q4 fraction = 0.54, 95% CI (0.39, 0.76), bootstrap P = 0.352, n = 57], so the confidence interval spanned the symmetric value of 0.5. Among the remaining pre-selected regions, the Q4 fraction ranged from a Q2-dominated 0.26 (posterior cingulate cortex) and 0.29 (dorsolateral prefrontal cortex) to a Q4-dominated 0.76 (dorsomedial prefrontal cortex), 0.80 (ventromedial prefrontal cortex), and 0.89 (angular gyrus). Thus, unlike the sensory regions, the PMC story-to-interruption transformation was not explained by a pure post-stimulus hemodynamic undershoot (Fig. S9).

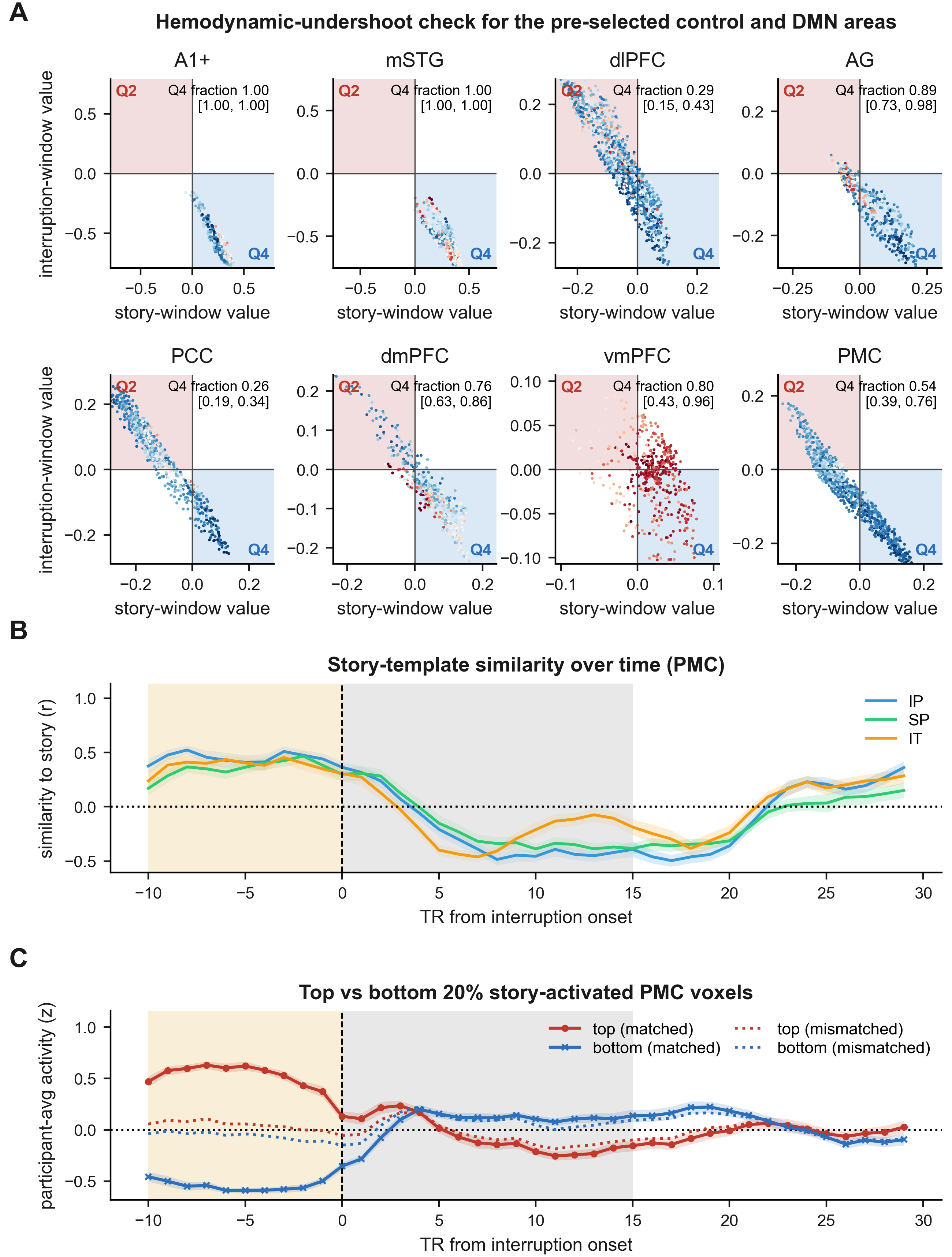

**Fig. S9.** Hemodynamic-undershoot control for the pre-selected regions. (A) For each of the eight pre-selected regions, we plot one dot for each voxel, comparing the grand-mean story-window value (x-axis) versus interruption-window value (y-axis). The grand means are pooled across epochs and participants (n = 57; 19 per interrupted condition). The “inversion” quadrants Q2 (story-negative to interruption-positive) and Q4 (story-positive to interruption-negative) are shaded red and blue respectively. The top-right text annotation reports the fraction of Q4 voxels among all inverting voxels (Q2 + Q4) with a 95% confidence interval generated by participant-wise bootstrapping. (B) Leave-one-participant-out similarity between each participant's PMC pattern and the group story-phase template, plotted over time from interruption onset (measured in TRs). Curves are shown for the three interruption conditions (n = 19 each); the similarity turns negative after onset and stays negative across the interruption window; it does not return toward zero after 10-20 s, as would be expected for a conventional hemodynamic undershoot. (C) BOLD time course of activated and deactivated voxels for each epoch. The 20% of voxels with the greatest (or least) story-phase BOLD signals are selected for each epoch and each participant, and their BOLD signal is then followed into the interruption phase. Solid lines depict BOLD signal from these voxels on the matching epoch (i.e. voxels selected and plotted for the same epoch) and dotted lines depict BOLD signal for mismatching epochs (i.e. voxels selected for activity on one epoch and then plotted based on activity in other epochs). The bottom voxels, when selected on the matching epoch, rise to sustained positive activity across the interruption, which cannot be explained by hemodynamic undershoot.

***S11. Story-to-interruption transformation: phase-wise z-score control***

Whole-run z-scoring could in principle contribute to the negative story-to-interruption relationship by centering the story and interruption phases relative to a common run-wide mean. We therefore repeated the transformation analysis after z-scoring each voxel separately across the story and interruption phases within each run. We excluded the first 5 TRs of each phase from the normalization process; all subsequent analysis steps were unchanged.

In the original whole-run-normalized analysis, the PMC story-to-interruption correlation was negative in all five comparisons [Fisher-z group mean = −0.245 (IP-IP), −0.209 (SP-SP), −0.171 (IT-IT), −0.139 (IP-IT), −0.163 (IT-IP); all sign-flip P < 0.001, n = 19 each]. After phase-wise z-scoring, the correlation remained negative in all five comparisons [IP-IP: −0.122, 95% CI (−0.151, −0.092); SP-SP: −0.073, 95% CI (−0.105, −0.040); IT-IT: −0.108, 95% CI (−0.145, −0.070); all sign-flip P ≤ 0.004, n = 19 each]. The two across-condition comparisons produced correlations of −0.103, 95% CI (−0.125, −0.081), P < 0.001 (IP-IT) and −0.076, 95% CI (−0.113, −0.040), P = 0.001 (IT-IP), n = 19 each.

The transformation also stayed epoch-specific in all three within-condition comparisons [IP-IP: Δ = −0.116, 95% CI (−0.143, −0.088); SP-SP: Δ = −0.073, 95% CI (−0.104, −0.042); IT-IT: Δ = −0.096, 95% CI (−0.130, −0.062); all P < 0.001, n = 19 each]. Thus, both the negative story-to-interruption relationship and its epoch specificity survived separate normalization of the story and interruption phases, indicating that the transformation was not created by whole-run normalization (Fig. S10).

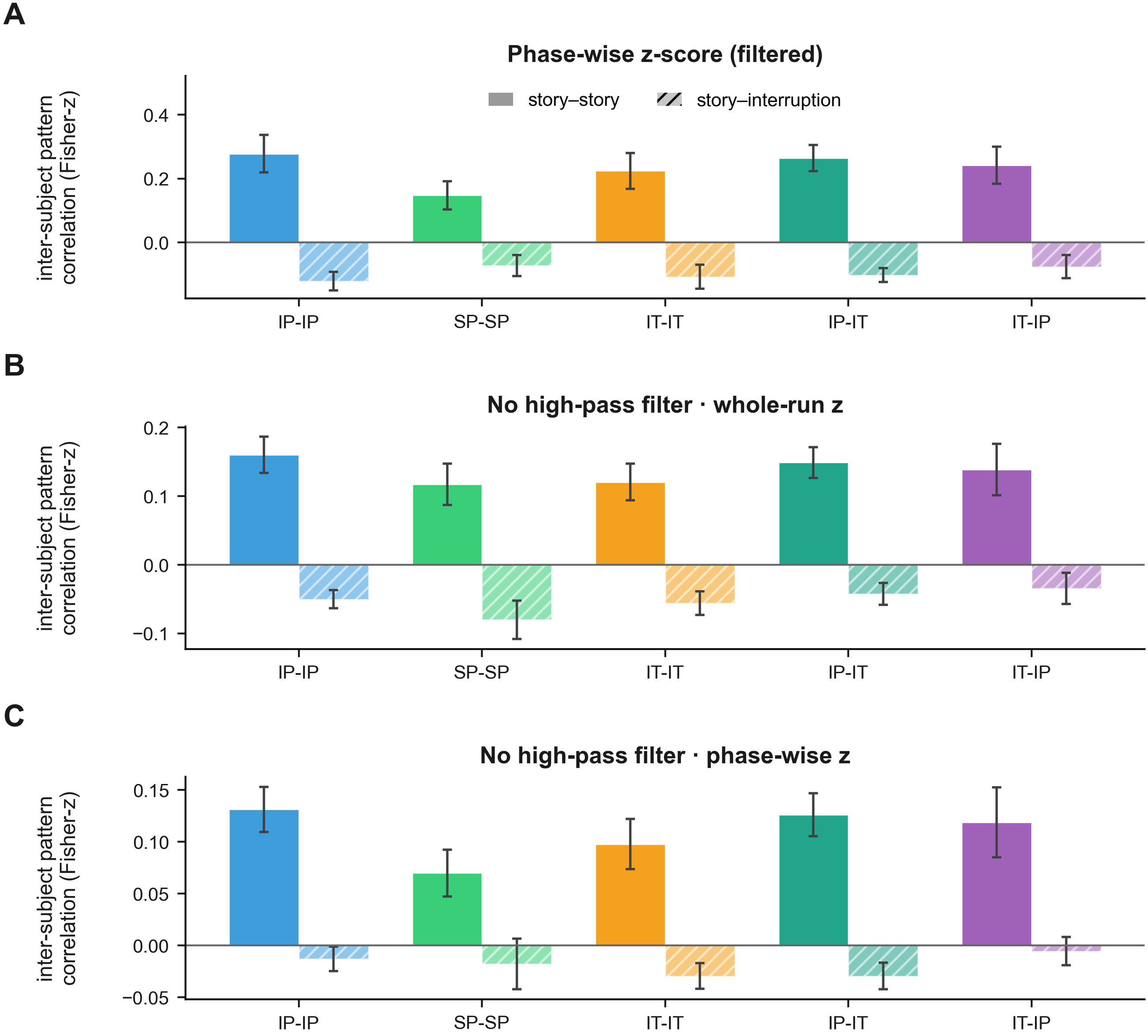

**Fig. S10.** PMC story-to-interruption transformation is robust to normalization and temporal high-pass filtering. Each panel shows story-story ISPC (solid) and story-to-interruption ISPC (hatched) within posterior medial cortex (Fisher-z group mean ± 95% confidence interval; n = 19 per comparison). The correlations are computed for the five comparisons (IP-IP, SP-SP, IT-IT, IP-IT, IT-IP). (A) Each voxel z-scored separately within the story and interruption phases, on the filtered main-pipeline data. (B) Unfiltered fMRI data (fMRIPrep minimal preprocessing pipeline, with no high-pass filter) with whole-run z-scoring. (C) Unfiltered fMRI data with phase-wise z-scoring. The negative story-to-interruption correlation (the inversion) persists under every normalization and filtering choice, except for the phase-wise SP-SP and IT-IP comparisons in (C), which do not reach significance.

***S12. Story-to-interruption transformation: high-pass-filter-off control***

We next tested whether the PMC transformation required temporal high-pass filtering, which could in principle contribute to anticorrelation between adjacent task phases. The PMC transformation analysis was repeated on data re-preprocessed with fMRIPrep and left temporally unfiltered (no temporal high-pass filter and no nuisance regression). The time series were extracted using both whole-run and phase-wise z-scoring. We also repeated the main-text interruption-pattern selectivity test on the same unfiltered data to determine whether the interruption pattern retained epoch-specific information without high-pass filtering.

With whole-run z-scoring, the PMC story-to-interruption correlation remained significantly negative in all five comparisons [Fisher-z group mean = −0.050 (IP-IP), 95% CI (−0.063, −0.037); −0.080 (SP-SP), 95% CI (−0.108, −0.052); −0.056 (IT-IT), 95% CI (−0.073, −0.039); −0.042 (IP-IT), 95% CI (−0.058, −0.027); −0.035 (IT-IP), 95% CI (−0.058, −0.012); all sign-flip P ≤ 0.004, n = 19 each]. Under the stricter combination of no high-pass filtering and phase-wise z-scoring, the negative correlation remained significant in three of the five comparisons: IP-IP [Fisher-z = −0.013, 95% CI (−0.025, −0.001), P = 0.035], IT-IT [−0.030, 95% CI (−0.042, −0.017), P = 0.004], and IP-IT [−0.029, 95% CI (−0.042, −0.017), P < 0.001; n = 19 each]. The SP-SP and IT-IP comparisons remained negative but did not reach significance [P = 0.147 and P = 0.217, n = 19 each].

The interruption-phase PMC pattern also remained epoch-specific in the IP condition without high-pass filtering. PMC patterns were more similar across participants for matching than mismatching epochs under both normalization procedures [whole-run z-scoring: selectivity = 0.016, 95% CI (0.010, 0.023), permutation P < 1 × 10−4; phase-wise z-scoring: selectivity = 0.016, 95% CI (0.010, 0.024), P < 1 × 10−4; n = 19 each]. Thus, high-pass filtering was not necessary for either the negative PMC story-to-interruption relationship or the epoch-specific interruption pattern. The transformation remained significant in all five comparisons with whole-run normalization and in three of five comparisons under the stricter combination of filter removal and phase-wise normalization (Fig. S10).

For comparison, A1+ showed a larger negative story-to-interruption correlation in the unfiltered whole-run data [Fisher-z = −0.134 (IP-IP), −0.148 (SP-SP)], consistent with its sensory-offset profile (Section S10). However, unlike PMC, the A1+ anticorrelation disappeared after phase-wise normalization (IP-IP = +0.012, SP-SP = +0.028; both not significant). Thus, the magnitude of the raw anticorrelation alone does not distinguish a representational transformation from a sensory-offset response; the phase-wise normalization and voxel-wise undershoot controls provide that distinction.

***S13. Story-to-interruption similarity time course without temporal filtering***

We next asked whether temporal high-pass filtering altered the time course of the PMC story-to-interruption anticorrelation. We recomputed the story-to-interruption similarity time course shown in Figure 3 using the unfiltered fMRI data (fMRIPrep minimal preprocessing, without high-pass filter) and compared it with the corresponding filtered time course around interruption onset.

Removing the high-pass filter and nuisance regressions decreased the absolute value of the anticorrelation but preserved the temporal profile of the correlation-by-time-lag function. The minimum was shallower in the fMRIprep data (story-to-interruption correlation approximately −0.05 to −0.07, compared with approximately −0.15 to −0.20 in the filtered data), but the two correlation-by-time-lag curves traced very similar shapes over time (r = 0.996 intact-pause, 0.994 scrambled-pause, 0.982 intact-ToM). The peak/trough timing was also preserved in the two pause conditions. Measuring from interruption onset, the correlation curve reached its minimum value at 18.0 s filtered and 19.5 s unfiltered in intact-pause, and at 19.5 s in both for scrambled-pause, that is, within one acquisition volume of each other; the story resumed no earlier than 22.5 s, so both minima fell inside the interruption. One exception was in the IT condition, where the trough of the curve moved (unfiltered: 10.5 s; filtered: 27.0 s), but this only reflected the fact that there were two very similar local minima: the filtered and unfiltered curves still showed a highly similar overall temporal profile (r = 0.98) and are visually extremely similar (Fig. S11).

Thus, although temporal high-pass filtering substantially increased the magnitude of the story-to-interruption anticorrelation, it had little effect on the temporal structure of the ISPC time course. If the anticorrelation were generated primarily by the high-pass filter, removing the filter should alter or abolish the temporal profile associated with the negative story-to-interruption relationship. Instead, the unfiltered data showed the same characteristic evolution of story-to-interruption similarity across the interruption, with nearly identical curve shapes and preserved trough timing in the two pause conditions. This indicates that filtering primarily amplified an underlying temporal relationship rather than creating it. Together with the persistence of a negative story-to-interruption relationship in the unfiltered data and the controls in Sections S10–S12, these results argue against a simple account in which the PMC anticorrelation is generated by the combination of post-stimulus undershoot and temporal high-pass filtering (Fig. S11).

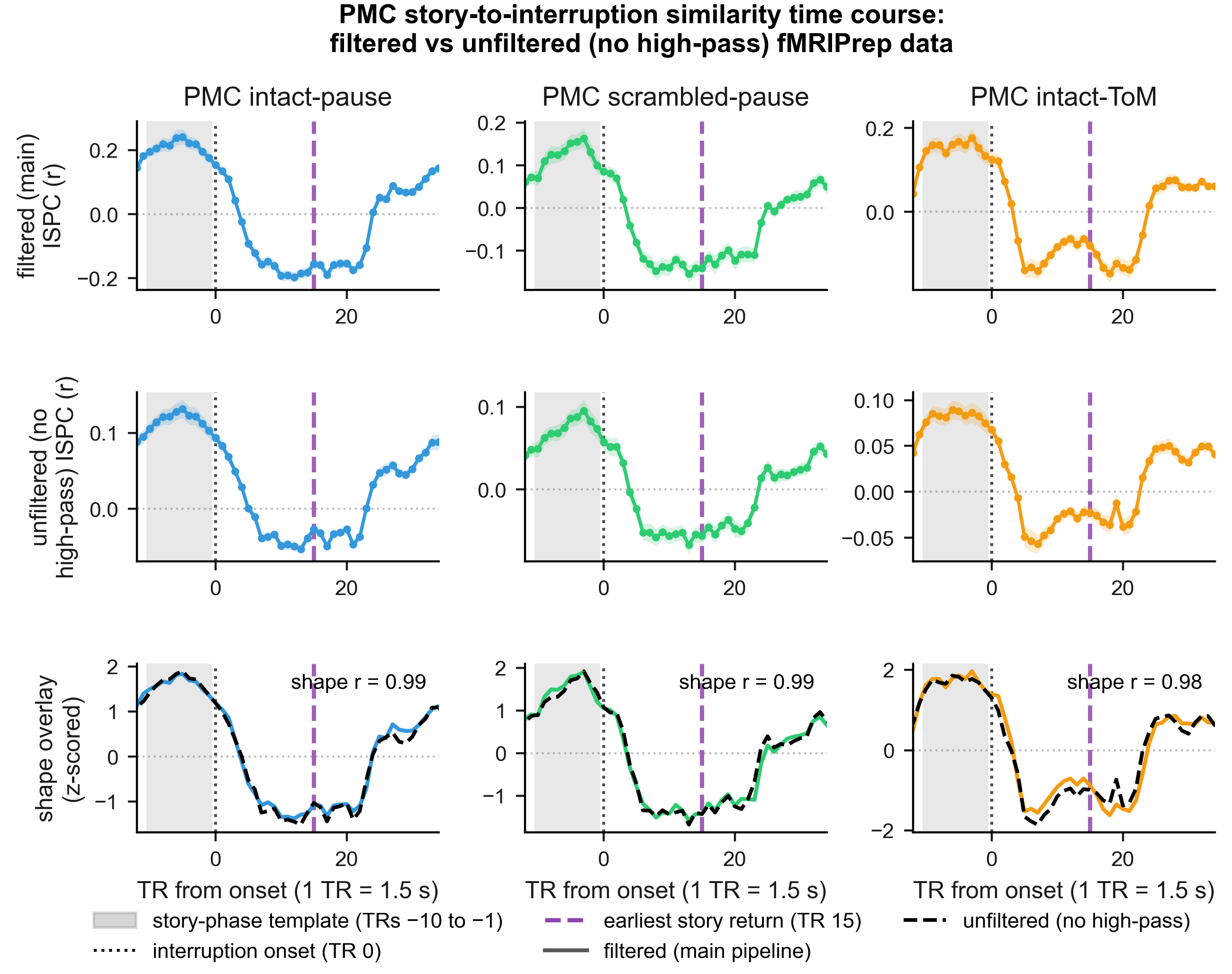

**Fig. S11.** PMC story-to-interruption similarity time course in filtered data (FSL pipeline) versus unfiltered data (fMRIPrep pipeline, lacking a high-pass filter). Row 1: filtered FSL-pipeline correlation-by-time-lag time course; Row 2: unfiltered fMRIPrep correlation-by-time-lag time course; Row 3: the two curves (z-scored) superimposed to compare shape only. In each panel the gray shaded area marks the time window for defining the story pattern template, the dotted vertical line marks interruption onset, and the purple dashed vertical line marks the earliest story return (TR 15). In the shape-overlay row, the solid curve is the filtered and the black dashed curve the unfiltered time course.

***S14. Story-to-interruption transformation: the within-participant flip predicts the shared interruption pattern***

We next asked whether the transformation measured across participants also held within participants, and whether it related to the strength of the shared interruption pattern persistence. For each participant, we computed an inversion index (the correlation between that participant's own PMC story-phase and interruption-phase patterns, averaged across the 17 epochs) and related it, across participants, to the strength of their shared interruption-phase pattern (each participant's mean ISPC with the other participants across epochs).

A stronger within-participant transformation (a more negative inversion index) was associated with a more reliably shared interruption-phase pattern in both intact-narrative conditions [IP: r = −0.73, 95% CI (−0.89, −0.59), Pearson P < 0.001; IT: r = −0.64, 95% CI (−0.87, −0.27), P = 0.003; n = 19 each]. The association was in the same direction under scrambling but did not reach significance [SP: r = −0.43, 95% bootstrap CI (−0.69, −0.12), Pearson P = 0.063, n = 19]. Thus, within the intact-narrative conditions, participants showing a stronger within-participant story-to-interruption transformation also showed a more persistently shared interruption-phase pattern.

***S15. Whole-brain Schaefer-400 story-to-interruption transformation***

We next asked how many cortical parcels exhibited a negative correlation between their story and interruption phase patterns similar to our pre-defined PMC area. The transformation test and the transformation-selectivity test were repeated at every parcel of the Schaefer 400-parcel atlas under the IP-IP, SP-SP, IT-IT, and IP-IT comparisons. For every parcel and epoch, each participant’s 10-TR story-phase template (ending one TR before interruption onset) was correlated with the comparison group’s interruption-phase template (the 10 TRs beginning 5 TRs after onset) at the corresponding epoch. We then Fisher-z transformed the resulting story-to-interruption correlation values and tested the group mean of the z values against zero. For this test, we used a one-sided sign-flip permutation test, because the prediction was in a single direction.

Across the 400 parcels, after Benjamini-Hochberg FDR correction, we observed significant negative story-to-interruption correlations in 139 parcels (IP-IP), 107 parcels (SP-SP), 29 parcels (IT-IT), and 58 parcels (IP-IT). The parcels exhibiting negative correlation were distributed across default-mode, salience/ventral-attention, frontoparietal-control, somatomotor and temporoparietal cortex rather than confined to any one network (Fig. S12). Thus, the negative story-to-interruption correlation was broadly distributed across cortex and was not unique to PMC.

However, parcels that passed the epoch-specific transformation-selectivity test (matching epochs more negatively correlated than mismatching) were far sparser. No parcel survived FDR correction in the IP-IP, SP-SP, or IP-IT. Thus, although the negative story-to-interruption transformation was widespread, its epoch-specific form was much more restricted at the whole-brain corrected threshold. Although no IP-IP parcel survived FDR correction, the strongest IP-IP selectivity values (the most negative matching-minus-mismatching differences) were concentrated in posteromedial cortex: seven of the eight most selective parcels among all 400 carried posterior-cingulate or precuneus labels (Fig. S12B).

Thus, even when the analysis was repeated across the whole cortex at the finer Schaefer-400 parcel scale, the strongest epoch-specific transformation effects converged on posteromedial cortex. This convergence supports the predefined PMC ROI and is consistent with a particular role for PMC in preserving story-context information in a transformed representational format.

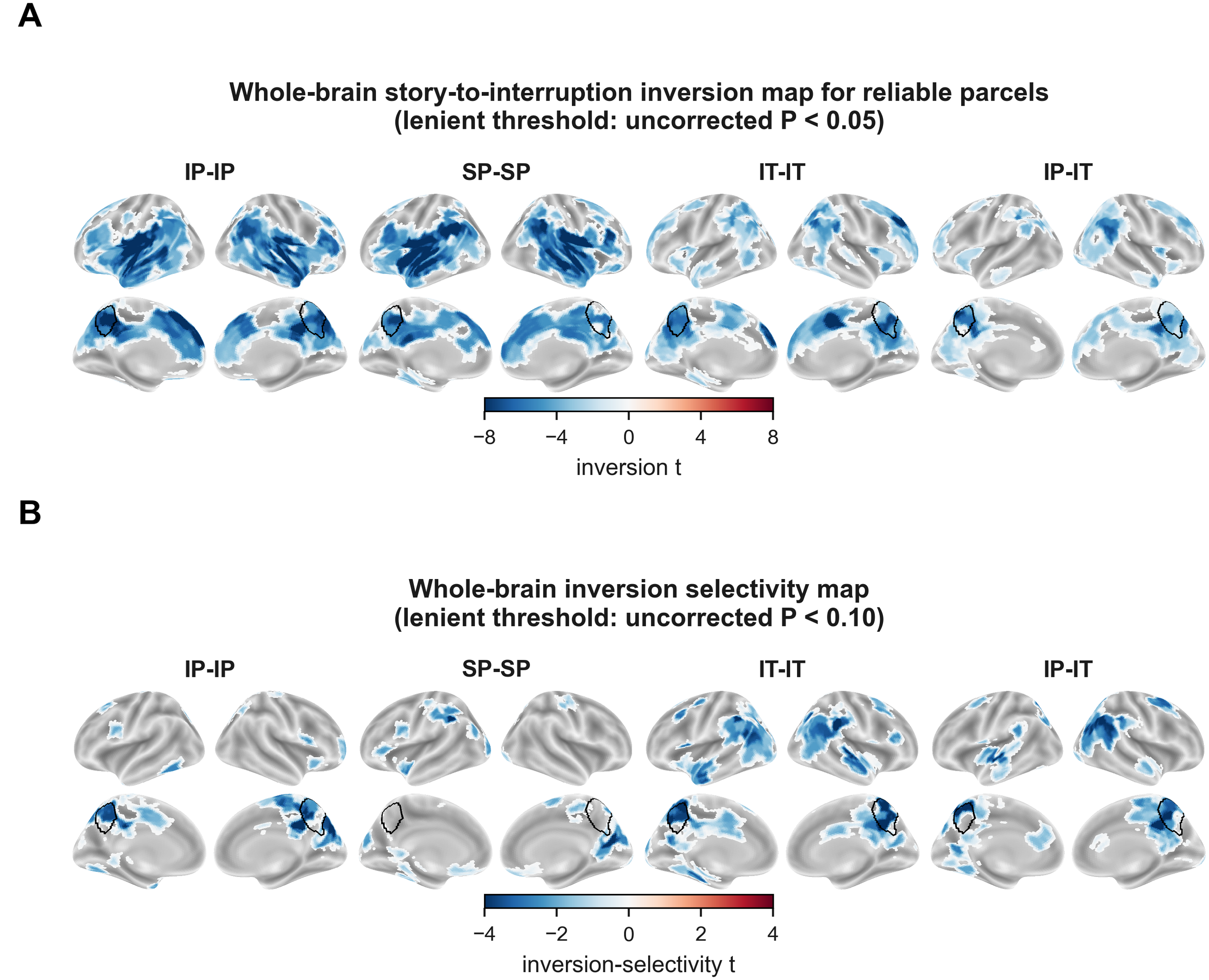

**Fig. S12.** Whole-brain maps of story-to-interruption correlations via the Schaefer-400 parcellation. (A) Inversion t-map (group-mean story-to-interruption correlation tested against zero; negative values indicate pattern reversal across the interruption) at every parcel that was reliable at an uncorrected sign-flip P < 0.05 in the condition supplying the interruption pattern and passed the transformation permutation test, for the IP-IP, SP-SP, IT-IT, and IP-IT comparisons (n = 19 per comparison). (B) Inversion-selectivity t-map, shown for parcels whose matching-versus-mismatching permutation reached an uncorrected *P* < 0.10 (with matching epochs more negatively correlated than mismatching). The perimeter of the posterior medial cortex (PMC) is outlined in black. Both panels deliberately use lenient display thresholds to show the spatial distribution of graded effects.

***S16. PMC pattern persistence measures a shared pattern that is stable across time***

In the main-text brain-behavior analyses, PMC pattern persistence was quantified as the mean ISPC across all pairs of TRs within the 10-TR interruption window. We asked whether this measure reflected a shared neural pattern that remained stable across the interruption period, rather than being driven primarily by temporally aligned pattern reliability across participants. In the corresponding interruption epochs’ 10 × 10 time-time ISPC block, diagonal cells (|i − j| = 0) compare patterns at the same TR across participants, whereas off-diagonal cells (|i − j| ≥1) compare patterns at different TRs. Thus, persistence of ISPC across increasingly distant off-diagonal cells provides a direct test of whether the shared interruption pattern remains stable across time.

We therefore recomputed PMC persistence using only cells with |i − j| ≥ L, where the minimum retained temporal lag L ranged from 1 (all off-diagonal cells) to 9 (only the two maximally separated corner cells), and refit the brain-behavior models at each lag (Fig. S13A) following the same procedure reported for the main-text.

The persistence effect was present in the off-diagonal (across-time) structure. For predicting neural resumption, the persistence slope coefficient was essentially unchanged after removing the diagonal from the persistence matrix. The association remained robust as progressively larger temporal separations were required [b = 0.016, t(53) = 4.87, P = 1.0 × 10^−5^ at |i − j| ≥ 1; b = 0.016, t(53) = 4.23, P = 9.3 × 10^−5^ at |i − j| ≥ 3; b = 0.013, t(53) = 2.85, P = 0.006 at |i − j| ≥ 6]. Only at the two-cell extreme did the predictive power of the PMC pattern decrease [b = 0.008, t(53) = 1.96, P = 0.056 at |i − j| ≥ 9].

For story recall, the slope coefficients from off-diagonal data were, if anything, slightly larger than the full-block estimate and they remained significant at every lag [b = 0.033, P = 0.004 at |i − j| ≥ 1; b = 0.036, P = 0.005 at |i − j| ≥ 3; b = 0.027, t(51) = 2.49, P = 0.016 at |i − j| ≥ 9]. Thus, the associations with both neural resumption and recall were supported by correlations between patterns measured at different time points, confirming that PMC pattern persistence captured a shared representation that remained stable across the interruption rather than merely same-TR pattern reliability across participants (Fig. S13).

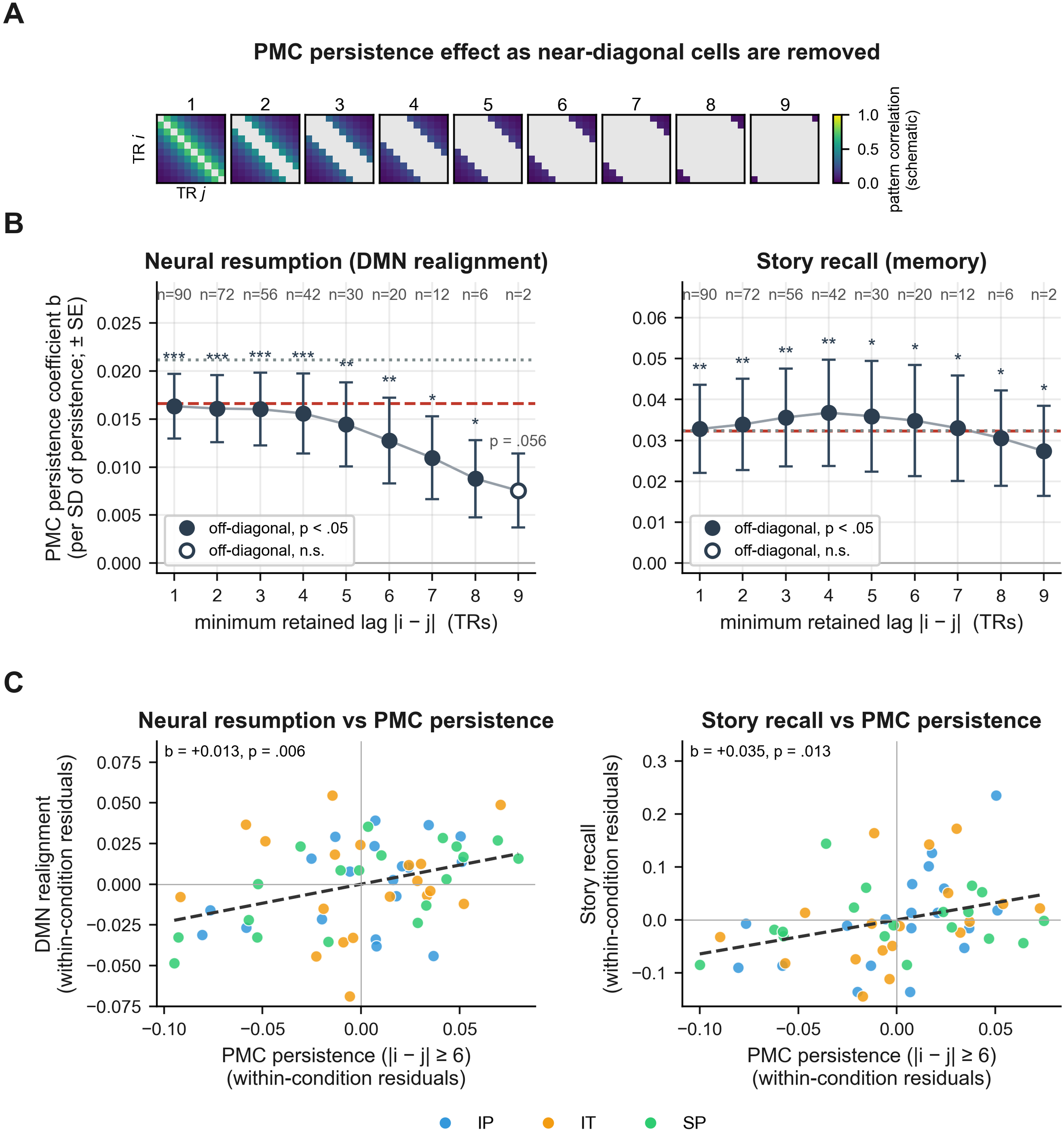

**Fig. S13.** Alternative method for measuring the temporal persistence of the PMC pattern during interruption. The effect survives removal of the matching-time ISPC cells, showing that the measure captures a shared PMC pattern that is stable across time rather than momentary same-TR pattern reliability. (A) Schematic of the 10×10 interruption time-by-time correlation block for each “minimum retained lag” |i − j|. Only the colored off-diagonal cells are averaged; gray matrix cells near the diagonal are excluded. (B) PMC persistence regression coefficient b (± standard error) as a function of the minimum retained time-lag, for neural resumption (default-mode-network realignment) and story recall. Lags at *P* < 0.05 are shown with filled circles and marked with asterisks directly above the top of each error bar (**P* < 0.05, ***P* < 0.01, ****P* < 0.001); the exact *P* value is printed only for the non-significant lags (open circles). The dashed red line indicates the full-block estimate without diagonal; the dotted gray line marks the diagonal-only estimate; the number of retained cells is printed above each column. (C) Scatter plots of PMC persistence computed from only the |i − j| ≥ 6 cells (20 of 100 block cells) against neural resumption (left) and story recall (right). The b printed in each panel is the regression coefficient for z-scored PMC persistence from the same model as panel (B), shown with its *P* value: it estimates the change in the outcome (neural resumption, or narrative recall) associated with a one-standard-deviation increase in PMC persistence. To adjust for condition, condition (IP, IT, SP) was entered as a categorical fixed effect in the regression; both plotted variables are shown as within-condition residuals, so the association reflects variation within conditions; dashed line shows the corresponding fit.

***S17. Comparison of story- and interruption-phase PMC pattern persistence***

As reported in the main text, persistence of the shared PMC interruption-phase pattern predicted default-mode-network pattern realignment at story resumption and later narrative recall; this PMC persistence explained variance in both outcomes beyond hippocampal boundary activity. To test whether the interruption-phase measure carried predictive information beyond a more general story-phase inter-subject pattern reliability effect, we computed the same persistence measure during the immediately preceding story phase. Specifically, for each interruption, story-phase persistence was computed from the 10 TRs immediately preceding interruption onset, excluding the onset TR. This story-phase measure was then entered into the same pooled ordinary-least-squares models used in the main text, either alone, together with interruption-phase persistence, or together with interruption-phase persistence and hippocampal boundary activity, for each outcome (DMN neural resumption and narrative recall).

The story-phase persistence predicted neural resumption when entered alone [b = 0.301, SE = 0.078, t(53) = 3.88, P < 0.001, 95% CI (0.145, 0.457)]. Story- and interruption-phase persistence were strongly correlated (r = 0.71). Once interruption-phase persistence was entered, the story-phase term no longer predicted resumption while the interruption-phase term did [story-phase b = 0.102, t(52) = 1.04, P = 0.305, 95% CI (−0.096, 0.300); interruption-phase b = 0.312, t(52) = 2.97, P = 0.005, 95% CI (0.101, 0.523)]. The same held with hippocampal boundary activity in the model [story-phase b = 0.047, t(51) = 0.48, P = 0.637; interruption-phase b = 0.318, t(51) = 3.15, P = 0.003; hippocampus b = 0.066, t(51) = 2.28, P = 0.027].

For narrative recall, the two persistence measures could not be statistically distinguished. The story-phase persistence predicted recall when entered alone [b = 0.694, SE = 0.242, t(51) = 2.87, P = 0.006, 95% CI (0.208, 1.18)]. When story- and interruption-phase persistence were entered together, however, neither retained a significant unique association with recall [story-phase b = 0.362, t(50) = 1.04, P = 0.303; interruption-phase b = 0.486, t(50) = 1.32, P = 0.193]. The same was true after hippocampal boundary activity was added [story-phase b = 0.221, t(49) = 0.63, P = 0.529; interruption-phase b = 0.501, t(49) = 1.39, P = 0.170; hippocampus b = 0.172, t(49) = 1.83, P = 0.073].

The two PMC measures were strongly correlated (r = 0.71), limiting our ability to distinguish their contributions to later narrative recall performance. Despite their strong association, interruption-phase ISPC predicted neural resumption beyond its story-phase counterpart, indicating that pattern persistence during interruption carries information relevant to post-interruption resumption that cannot be solely explained by its preceding story-phase pattern reliability.
